# Dimensionality reduction of calcium-imaged neuronal population activity

**DOI:** 10.1101/2022.03.11.480682

**Authors:** Tze Hui Koh, William E. Bishop, Takashi Kawashima, Brian B. Jeon, Ranjani Srinivasan, Sandra J. Kuhlman, Misha B. Ahrens, Steven M. Chase, Byron M. Yu

## Abstract

Calcium imaging has been widely adopted for its ability to record from large neuronal populations. To summarize the time course of neural activity, dimensionality reduction methods, which have been applied extensively to population spiking activity, may be particularly useful. However, it is unclear if the dimensionality reduction methods applied to spiking activity are appropriate for calcium imaging. We thus carried out a systematic study of design choices based on standard dimensionality reduction methods. We also developed a novel method to perform deconvolution and dimensionality reduction simultaneously (termed CILDS). CILDS most accurately recovered the single-trial, low-dimensional time courses from calcium imaging that would have been recovered from spiking activity. CILDS also outperformed the other methods on calcium imaging recordings from larval zebrafish and mice. More broadly, this study represents a foundation for summarizing calcium imaging recordings of large neuronal populations using dimensionality reduction in diverse experimental settings.

## Introduction

Computations in the brain occur through the coordinated, time-varying activity of populations of neurons. Dimensionality reduction is a class of statistical methods commonly used for summarizing neural population activity^1–3^. It transforms high-dimensional neural recordings, such as spiking activity from a population of recorded neurons, into compact low-dimensional representations termed *latent variables*. These low-dimensional representations facilitate the investigation of how neural population activity varies over time, across experimental conditions, and across repeated experimental trials of the same condition. In particular, dimensionality reduction has been used to uncover neural mechanisms underlying decision making^4^, motor control^5^, learning^6^, working memory^7^, sensorimotor timing^8^, attention^9^, olfaction^10^, speech^11^, and more.

Dimensionality reduction has typically been applied to electrophysiological recordings. In the last decade, optical imaging has been widely adopted to record from large populations of neurons given its ability to sample neurons densely within the field of view, track neurons over long periods of time, and label neurons by cell type or projection, among other advantages^12^. A leading type of optical imaging is calcium imaging, which uses calcium indicators to track the transient increase in intracellular calcium levels that accompanies electrical spiking activity^13^. These changes in calcium levels are then optically recorded via changes in fluorescence. Calcium imaging has the capability of imaging even the whole brain of some small animals (e.g., larval zebrafish) at single neuron resolution^14^, albeit at a lower temporal resolution than electrical recordings.

With the increasing use of calcium imaging, many studies are now beginning to apply dimensionality reduction to calcium imaging recordings^15–26^. A critical question is whether the same dimensionality reduction methods applied to study spiking activity are also appropriate for calcium imaging recordings^27^. Here, we focus on single-trial time courses of latent variables, termed *neural trajectories*. This enables the study of how trial-to-trial differences in the time course of neural activity relate to trial-to-trial differences in perception, decision making, and behavior^8,16,28–31^. We seek to understand if the neural trajectories extracted from calcium imaging recordings match the neural trajectories that would have been extracted from the spiking activity underlying these recordings. A key reason why the neural trajectories would be different is the slow, indicator-dependent decay of measured calcium levels after each spiking event^13^. This decay introduces temporal correlations in the calcium imaging recordings that would not be present with spiking activity alone. Deconvolution is a technique that aims to recover spiking activity from calcium imaging recordings^32^. However, deconvolution techniques do not, as yet, recover the underlying spikes exactly^33^. Thus, the neural trajectories that are extracted from calcium imaged activity may be quite different from the neural trajectories extracted from spiking activity (Fig. 1), which may limit the use of calcium imaging for studying population activity time courses. In this work, our central goals are to i) systematically study the appropriateness of dimensionality reduction methods for summarizing the time course of calcium imaging recordings, and ii) propose a new dimensionality reduction method that is tailored for extracting neural trajectories from calcium imaging recordings.

**Figure 1.**
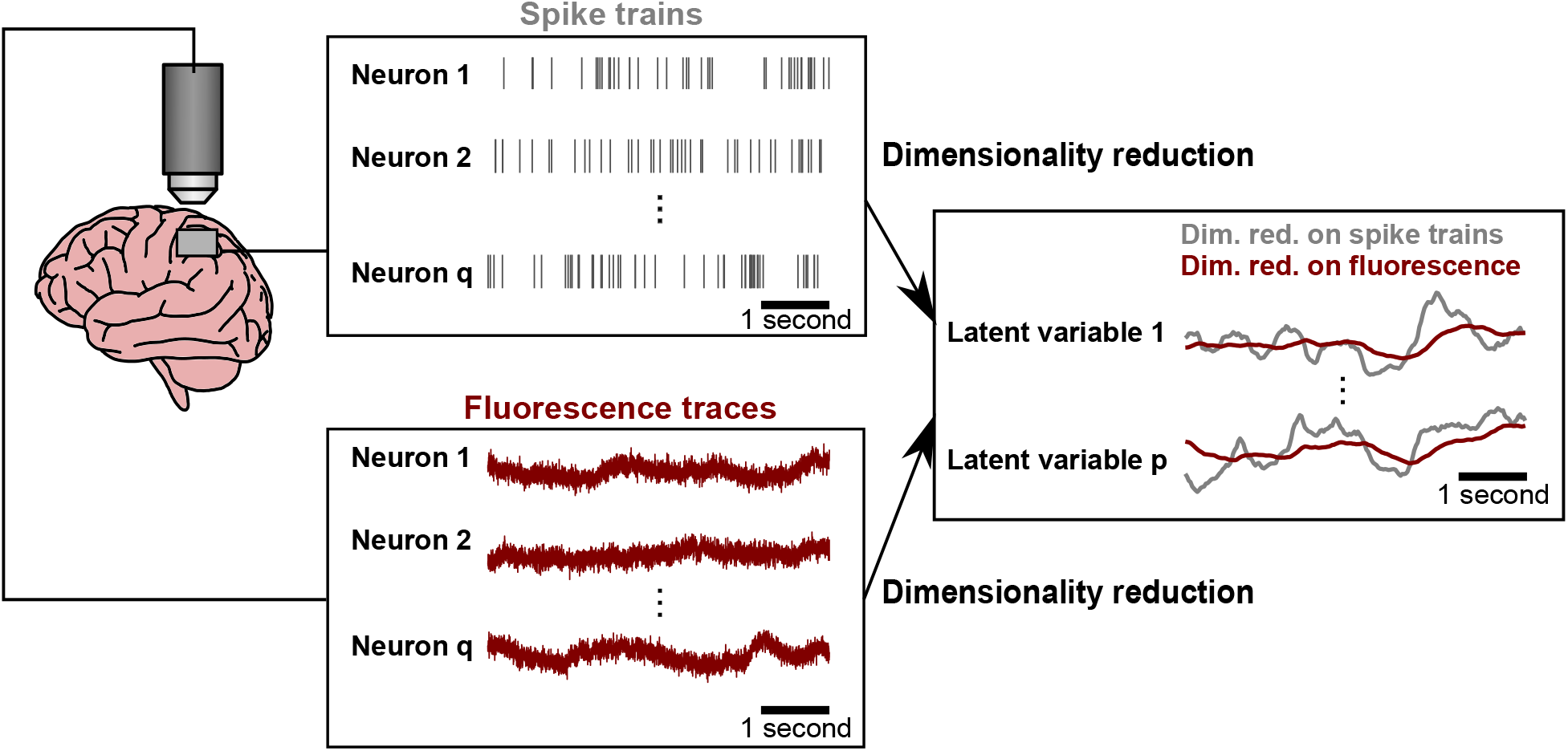
Dimensionality reduction of neuronal population activity based on electrophysiological recordings versus calcium imaging can yield different low-dimensional representations. Left, spike trains (grey) and fluorescence traces (maroon) recorded from the same hypothetical population of neurons. Right, different latent trajectories resulting from dimensionality reduction applied to spike trains (grey) versus fluorescence traces (maroon).

We sought to address three questions. First, we asked if deconvolution should be used with dimensionality reduction when extracting neural trajectories, and if so, how it should be applied. Second, we asked how different experimental variables (e.g., the decay constant of the calcium indicator, the timescale of the latent time courses, and the number of imaged neurons) impact the ability to recover neural trajectories from calcium imaging recordings. Third, we asked if it is necessary for the dimensionality reduction method to employ a dynamics model for the latent variables, as such a model might enable the time course of the neural trajectory to be more cleanly separated from the time course of calcium decay.

We addressed these questions by comparing several approaches, including i) standard dimensionality reduction applied directly to the recorded fluorescence; ii) a two stage-method in which deconvolution is applied separately to each neuron’s fluorescence trace to estimate spiking activity, then standard dimensionality reduction is applied to the estimated spiking activity; and iii) a novel unified method that we propose here (Calcium Imaging Linear Dynamical System, CILDS), which performs deconvolution and dimensionality reduction jointly. We first applied these methods to simulated fluorescence traces, in which we systematically varied several experimental variables over a wide range. We then applied these methods to calcium imaging recordings from the dorsal raphe nucleus of larval zebrafish and the primary visual cortex of mice. Across these settings, we found that CILDS outperformed the other methods. This was especially true when the neural activity fluctuated more quickly over time (timescale of tens to hundreds of milliseconds). We also found that, to accurately peer through the calcium decay to obtain accurate neural trajectories, it is necessary to include a latent dynamical model, as in CILDS. Overall, our work provides a foundation for using dimensionality reduction to summarize the time course of calcium imaging recordings.

## Results

Dimensionality reduction is typically applied to spike trains recorded from a population of neurons, yielding a low-dimensional representation of that activity. Calcium imaging provides a transformed view of those spike trains. Our central goal is to develop dimensionality reduction methods appropriate for calcium imaging recordings to recover the same low-dimensional representation as that obtained from spike trains. To do so, we systematically compare three approaches.

For the first approach, we applied a standard dimensionality reduction method directly to the recorded fluorescence traces from calcium imaging (Fig. 2a, top). Here we chose to use a latent Linear Dynamical System (LDS), which is among the most basic methods for extracting neural trajectories. Conceptually, an LDS seeks to explain the temporal structure in the data using latent variables that vary smoothly over time. We will examine the necessity of using a latent dynamical model, like LDS, in a later section.

**Figure 2.**
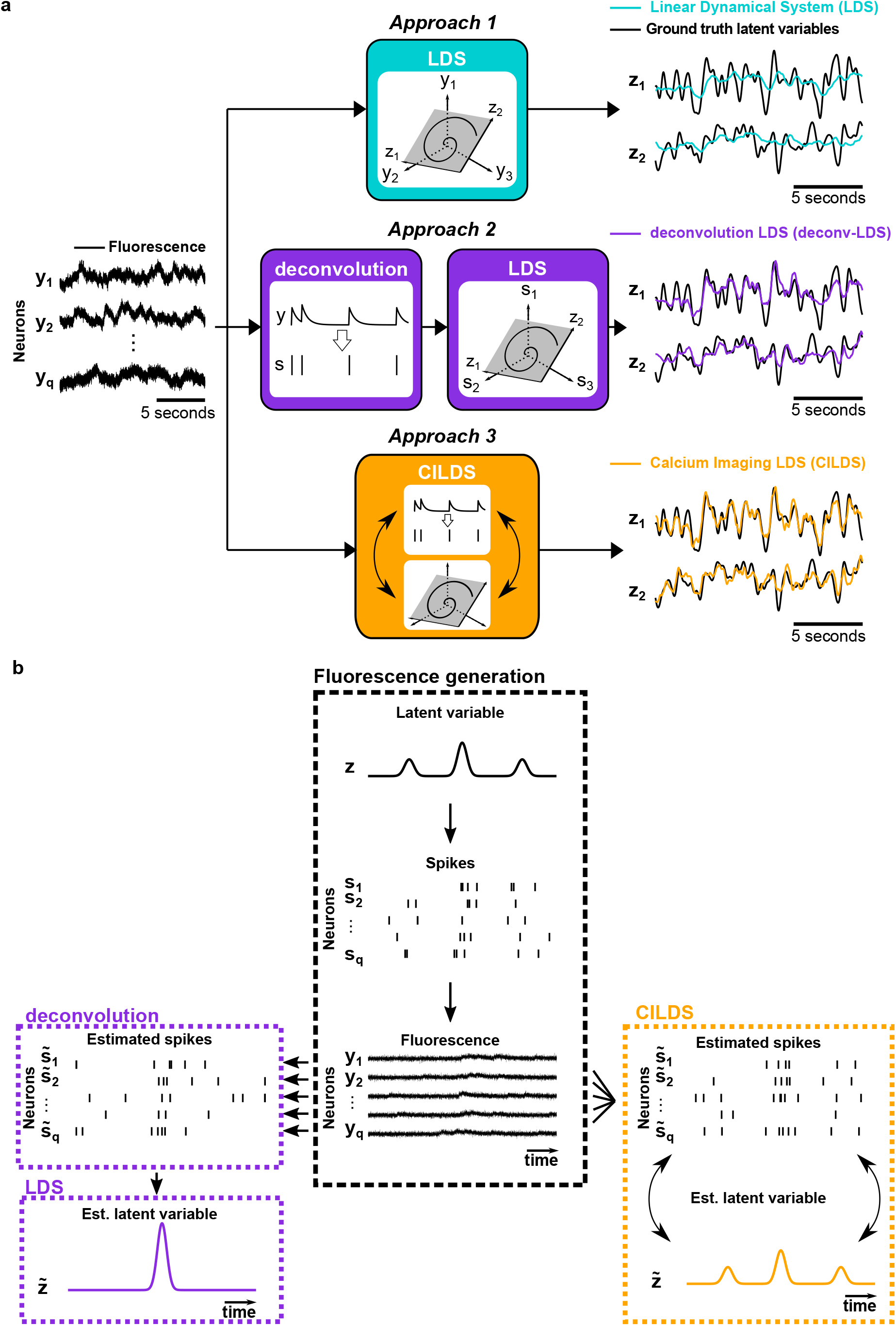
Comparison of three classes of dimensionality reduction methods. **(a)** Each of the three classes of methods was applied to the simultaneously-recorded fluorescence of a population of neurons (*y*_1_, *y*_2_, …, *y*_*q*_) to extract latent variables. Top, Approach 1: a standard dimensionality reduction method (e.g., LDS) applied directly to calcium imaging recordings, extracting corresponding low-dimensional latent variables at each time point (illustrated here with two dimensions, *z*_1_ and *z*_2_). Middle, Approach 2: deconvolution is applied separately to each neuron’s fluorescence trace to estimate its underlying spiking activity (*s*_1_, *s*_2_, …, *s*_*q*_). A standard dimensionality reduction method (e.g., LDS) is then applied to the estimated spiking activity to extract latent variables (*z*_1_ and *z*_2_). Bottom, Approach 3: A unified method (e.g., CILDS) that takes calcium imaging recordings as input and performs deconvolution and dimensionality reduction simultaneously to extract the latent variables (*z*_1_ and *z*_2_). **(b)** Cartoon depicting the intuition behind the difference between Approaches 2 and 3. Center column: a latent variable *z* (representing, for example, common input) is used to generate spike trains which, in turn, are used to generate fluorescence traces. Left column: if deconvolution is performed neuron by neuron (Approach 2, deconv-LDS), it is unable to leverage the shared activity fluctuations among neurons to dissociate the calcium transients from the underlying shared spiking activity (i.e., the estimated latent variable). Right column: A unified method (Approach 3, CILDS) is applied to all neurons together and is therefore better able to dissociate the calcium transients from the underlying shared spiking activity among neurons (i.e., the estimated latent variable). As a result, CILDS more accurately estimates the ground truth latent variable than deconv-LDS. Note that the estimated spiking activity is depicted here as spike trains for visual clarity, even though they are in fact continuous-valued time courses.

Each time a neuron spikes, intracellular free calcium increases, then decays slowly over time. The calcium indicator kinetics influence the measured decay time, resulting in fluorescence traces whose intensity decays over hundreds of milliseconds to seconds, depending on the particular calcium indicator used^13^. This decay transient induces temporal correlations in the fluorescence measurements which are input to the LDS, which might attempt to capture these correlations in its latent variable estimates. This motivates our second approach, which first deconvolves each fluorescence trace separately, and then applies a LDS to the resulting estimated spiking activity. The deconvolution serves to remove a substantial portion of the calcium decay transient, producing activity traces similar to spike trains (or time varying firing rates). We term this two-stage method *deconv-LDS* (Fig. 2a, middle). Here we deconvolved the fluorescence traces with OASIS^32,34^, a deconvolution method that has been widely used in calcium imaging studies^20,23,24,35^.

In deconv-LDS, each neuron is deconvolved independently, and the stages of deconvolution and dimensionality reduction are performed sequentially. We asked if performing these stages jointly would lead to more accurate recovery of the latent variables (Fig. 2a, bottom). More specifically, we hypothesized that a sequential method (e.g., deconv-LDS) may inadvertently discard some of the shared activity amongst neurons due to the independent deconvolution of each neuron (Fig. 2b, left). Since the latent variables are intended to capture the shared spiking activity among neurons (and not the calcium decay, which is independent across neurons), it might be possible to better separate the calcium decay from the latent dynamics by considering all the neurons together, and performing the two stages of deconvolution and dimensionality reduction jointly. This allows the dimensionality reduction component to influence the deconvolution estimates, and vice versa (Fig. 2b, right). Thus, for our third method we developed a unified approach, *CILDS*, in which dimensionality reduction and deconvolution are performed jointly (Fig. 2a, bottom).

### Should deconvolution be used with dimensionality reduction, and if so, how?

One might postulate that first deconvolving fluorescence to estimate spiking activity would be beneficial for recovering the same underlying latent variables that one would have recovered from the spikes themselves^27^. Indeed, multiple studies have applied dimensionality reduction to deconvolved spiking estimates^19–24,35^. However, deconvolution is subject to particular statistical modeling assumptions (as is any statistical method) and usually does not recover the underlying spikes exactly. Therefore it is unclear how, or even if, deconvolution should be used with dimensionality reduction. We addressed this question by comparing the three approaches described above (Fig. 2a).

With calcium imaging recordings, we typically do not have simultaneous electrophysiological recordings from the same neurons, and thus the “ground truth” latent variables are unknown. To directly compare each method’s ability to extract latent variables, we designed a simulation framework in which we created known ground truth latent variables with smoothly-varying time courses. These latent variables were used to generate spike trains which, in turn, were used to generate fluorescence traces (see Methods and Supplementary Fig. 1). We then applied each of the three approaches to these simulated fluorescence traces to assess how accurately they recovered the ground truth latent variables. Examples of two combinations of experimental variables are illustrated in Fig. 3a.

**Figure 3.**
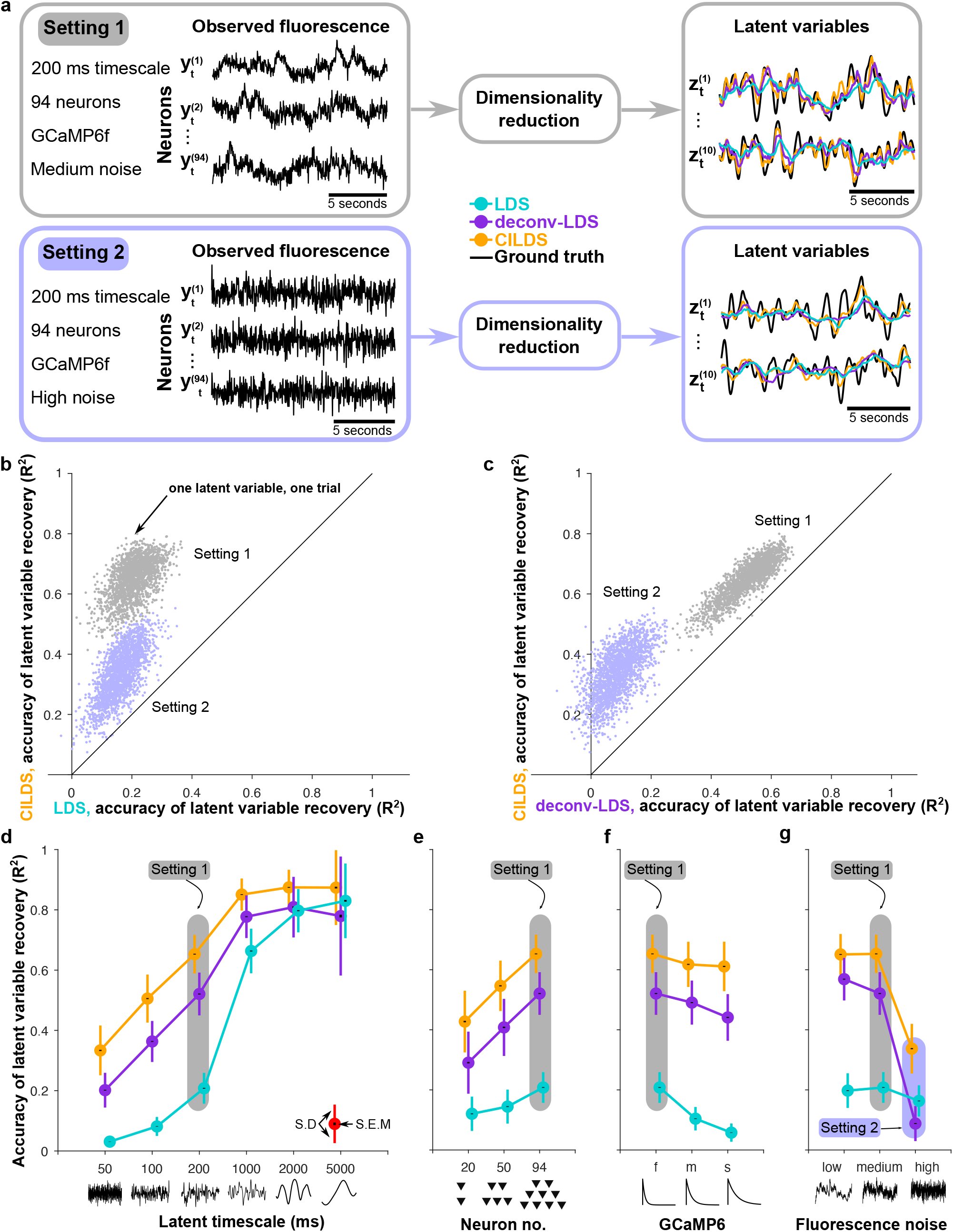
CILDS more accurately recovers the ground truth latent variables than the other two approaches in simulation. **(a)** Example simulated fluorescence traces (left panels) and estimated latent variables (right panels) for two combinations of experimental variables. Setting 1 corresponds to a latent timescale of 200ms, 94 neurons, calcium decay corresponding to GCaMP6f, and medium fluorescence noise (see Methods). Setting 2 is the same as Setting 1, but with high fluorescence noise. Each of the three dimensionality reduction approaches introduced in Fig. 2 (LDS, cyan; deconv-LDS, purple; CILDS, orange) is applied to the simulated fluorescence traces. The latent variables extracted by each method can then be compared to the ground truth latent variables (black). **(b-c)** Accuracy of latent variables estimated by CILDS versus that of (b) LDS and (c) deconv-LDS. Accuracy is measured by the *R*^2^ between the estimated and ground truth latent variables. Each point represents one latent variable on one trial. **(d-g)** Accuracy of latent variable recovery, as the (d) latent timescale, (e) number of neurons, (f) GCaMP6 indicator decay time constant, and (g) fluorescence noise level was varied. In each panel (d-g), one of the experimental variables was varied, while the other three variables were held constant at the Setting 1 values. The common point across the four panels is Setting 1 (shaded gray). Setting 2 (shaded purple) only appears in panel (g) because panels (d)-(f) correspond to medium rather than high fluorescence noise. The *R*^2^ for other combinations of experimental variables are shown in Supplemental Fig. 2. Colored error bars indicate standard deviation, and black error bars indicate standard error across n=2000 latent variables (see Methods). The points are horizontally offset for visual clarity.

We found that in both settings, CILDS outperformed the other two methods, returning more accurate estimates of the ground truth latent variables (Fig. 3b-c, points above the diagonal). By operating on the entire population of neurons together, CILDS was better able to separate the calcium transients from the shared activity among neurons (i.e., the latent variables) compared to deconv-LDS, which deconvolves the activity of each neuron individually, and LDS, which makes no attempt at this separation. Taken together, when extracting single-trial neural trajectories, one should use deconvolution (Fig. 3b) jointly with dimensionality reduction, as in CILDS (Fig. 3c).

### How do different experimental variables impact the ability to recover latent activity?

We next asked how different experimental variables affect the accuracy of the recovered latent variables. We designed the simulation framework to enable us to systematically vary experimentally relevant variables. These variables comprise four axes along which we can explore different combinations of values that might mimic a particular experimental paradigm. We varied the timescale of the latent time courses from 50ms to 5000ms, the number of neurons from 20 to around 100, the calcium decay timescales to match GCaMP6f, GCaMP6m, and GCaMP6s (fast, medium, slow)^13^, and the variance of the added imaging noise that was independent of the calcium and spiking activity (Supplementary Table 1). We then assessed how the accuracy of the dimensionality reduction methods changed as we systematically changed these variables.

As we increased the timescale of the latent variables, all dimensionality reduction methods improved their accuracy in estimating the ground truth latent variables (Fig. 3d). This occurs because with slower latent fluctuations, the latent variables become less independent across time. As a result, all methods can leverage future and past time points to better estimate the latent variables at the current time point. When the latent timescale is slow (order of seconds), the calcium indicator decay is shorter relative to the latent timescale, and influences the neural activity less. Therefore a method that is not able to disambiguate between the time course of the latent variables and the calcium decay (e.g., LDS) can still accurately recover the latent variables (see latent timescale of 5000 ms in Fig. 3d). However, at faster latent timescales (tens to hundreds of milliseconds) which reflect the timescales of many sensory, cognitive, and motor functions^35,36^, it is critical to use a method that accounts for the calcium decay, as both CILDS and deconv-LDS do (see latent timescale of 50 ms in Fig. 3d).

Next, when we increased the number of “recorded” neurons, all three methods improved in their ability to reconstruct the ground truth latent variables (Fig. 3e). This makes sense because each neuron provides a different, noisy view of the underlying latent variables. With more neurons, all methods are better able to “triangulate” the values of the latent variables. Across the entire range of neurons tested, CILDS outperformed deconv-LDS, which outperformed LDS (Fig. 3e).

We also varied the time constant of the GCaMP calcium indicator decay to match GCaMP6f, 6m, and 6s (from fast to slow) (Fig. 3f). All methods performed worse as the decay time constant increased. This occurred because the slower the calcium indicator is, the less the resulting fluorescence signal resembles the original spike train, which increases the difficulty in disambiguating between the latent time course and the calcium decay (Fig. 3f).

Finally, we varied the amount of noise added to the fluorescence, which reflects imaging noise independent of calcium and spiking activity (Fig. 3g). As the variance of the noise increased, all three methods performed worse as expected, although the extent of the performance degradation differed across methods. As the fluorescence noise variance increased, CILDS continued to outperform the other two methods (Fig. 3g). This indicates that leveraging the population of neurons for simultaneous deconvolution and dimensionality reduction, as done by CILDS, provides statistical power to mitigate a loss in accuracy due to increased fluorescence noise (Fig. 2b).

Overall, CILDS performed as well or better than the other two methods in every simulated setting we tested (Fig. 3d-g, orange higher than purple and cyan, also see Supplementary Fig. 2 for additional combinations of simulation parameter settings). We additionally found that deconv-LDS usually outperformed LDS in accuracy of recovered latent variables, consistent with Wei et al., which applied PCA to trial-averaged activity^27^ (Fig. 3d-g, purple higher than cyan). Using deconvolution is particularly important in regimes where the time scales of neural activity (i.e., the latent timescales) are faster than that of the calcium decay, which is the case for many commonly-studied brain functions.

### Is it necessary to include a latent dynamical model in the dimensionality reduction method?

All three dimensionality reduction methods considered so far explicitly attempt to extract latent variables that evolve smoothly over time via a latent dynamical model. We asked if the latent dynamical model was necessary for separating the calcium decay from the neural trajectories. To address this, we developed a method called Calcium Imaging Factor Analysis (CIFA, see Methods). CIFA is identical to CILDS, except that CIFA’s latent variables are independent across time (i.e., there is no dynamical model) by analogy to conventional factor analysis.

We then compared the performance of CIFA to CILDS. We found that CILDS, which uses a latent dynamical model, recovered the latent variables more accurately than CIFA, which does not use a latent dynamical model (Fig. 4a; additional cases are shown in Supplementary Fig. 3). As the latent timescale increased, CILDS more accurately recovered the latent variables (Fig. 4a, also shown in Fig. 3d). Correspondingly, CILDS correctly ascertained that the increased smoothness in the fluorescence was due to a longer latent timescale and not a longer calcium decay time constant (Fig. 4b, Supplementary Fig. 3). In contrast, as the latent timescale increased, CIFA recovered the latent variables less accurately (Fig. 4a). This occurs because CIFA erroneously attributed the increased smoothness in the fluorescence to a slower calcium decay (Fig. 4b, Supplementary Fig. 3), rather than a longer latent timescale. Unlike CILDS which can attribute temporal smoothness in the fluorescence to two possible sources (latent variables that evolve smoothly over time and calcium decay), CIFA can attribute temporal smoothness to only one possible source (calcium decay). This indicates that using a latent dynamical model is necessary for peering through the calcium decay transients to accurately recover the neural trajectories.

**Figure 4.**
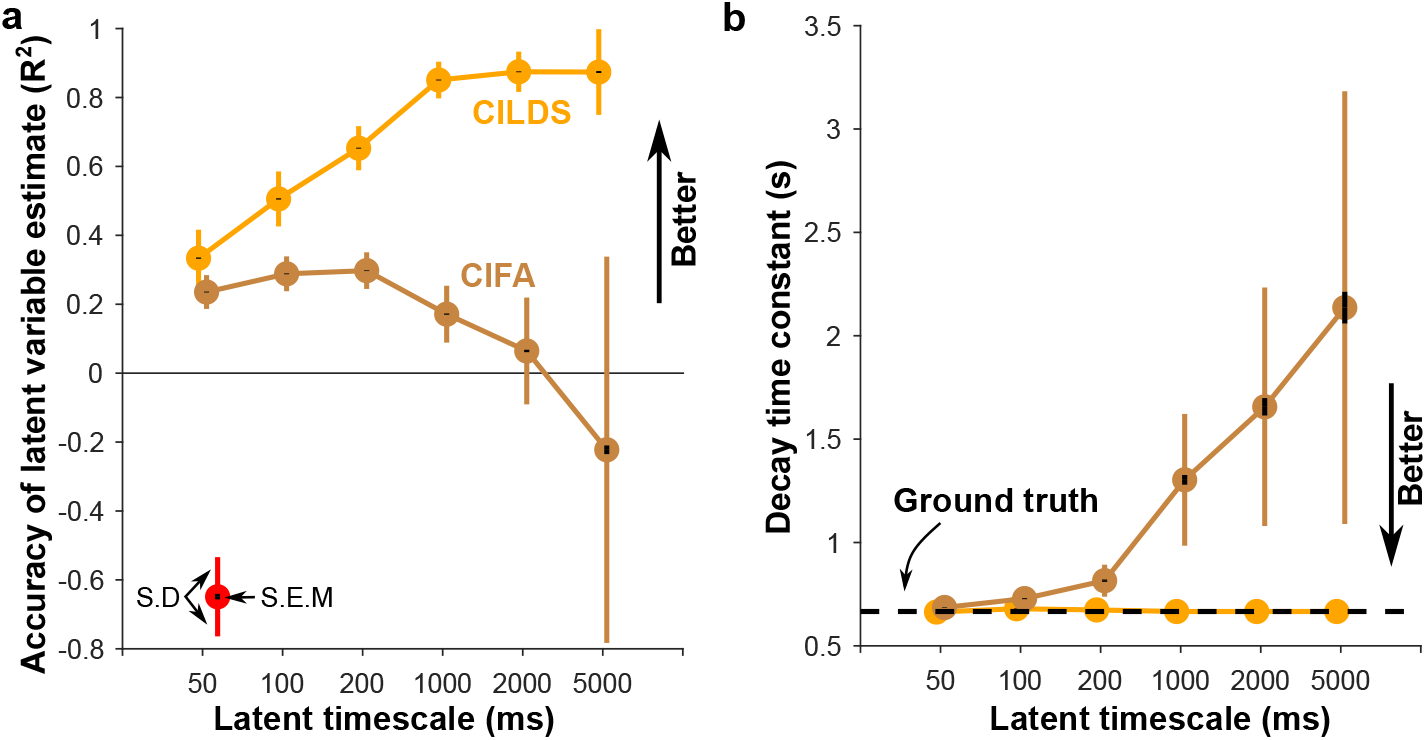
A latent dynamical model is necessary for accurately recovering neural trajectories. Comparison of a method that has no latent dynamics (Calcium Imaging Factor Analysis, CIFA) to CILDS using simulated fluorescence traces. The simulation parameters are GCaMP6f with 94 neurons and medium noise, as in Fig. 3d. **(a)** As the latent timescale increases, the ability of CILDS (orange) to accurately recover the neural trajectories increases, whereas that for CIFA (brown) decreases. Note that the *R*^2^ can be < 0 because these results are cross-validated. The CILDS curve shown here is the same as in Fig. 3d. **(b)** Calcium decay time constant estimated using CIFA (brown) and CILDS (orange) for different simulated latent timescales. Dashed black line indicates ground truth decay time constant. CILDS accurately estimates the decay time constant across all latent timescales tested, whereas CIFA overestimates the decay time constant as the latent timescale increases. Coloured error bars indicate standard deviation, and black error bars indicate standard error across *n* = 2000 latent variables (see Methods).

### CILDS outperforms other methods on calcium imaging recordings

To assess whether the advantages of CILDS also hold in real data, we applied each of the dimensionality reduction methods described above (CILDS, deconv-LDS, LDS, and CIFA) to calcium imaging recordings in two experimental contexts: larval zebrafish and mice. To emphasize the generality of our findings, these two experimental settings involve not only different animal species, but also different brain areas, behavioral tasks, and properties of the recorded fluorescence (see below). In these experiments, the spiking activity of the neurons was not recorded, and thus the ground truth latent variables are unknown. To quantify the accuracy of each method, we adopted a leave-neuron-out procedure used in previous studies^37,38^, where we estimate the latent variables using all-but-one neuron, and then assess how well these latent variables predict the recorded fluorescence of the held-out neuron (see Methods). A more accurate prediction of the fluorescence of the held-out neuron indicates that the latent variables are a better summary of the population activity.

The first experimental context involves larval zebrafish engaged in a “fictive swimming” motosensory gain adaptation task (Fig. 5a)^39^. Calcium imaging was performed on neurons expressing GCaMP6f in dorsal raphe nucleus (DRN) using light-sheet microscopy in three fish (19 to 22 neurons; mean: 20) (Fig. 5a). We applied each dimensionality reduction method to these recordings and assessed their performance using the leave-neuron-out prediction procedure (Fig. 5b). We found that CILDS more accurately predicted the fluorescence of the held-out neurons than the other methods, as quantified by the correlation between the predicted and recorded fluorescence. CILDS outperformed LDS (Fig. 5c; 63% of neurons above the diagonal), deconv-LDS (Fig. 5d; 85% of neurons above the diagonal), and CIFA (Fig. 5e; 73% points above the diagonal). These results were also true for each fish individually (Supplementary Fig. 5).

**Figure 5.**
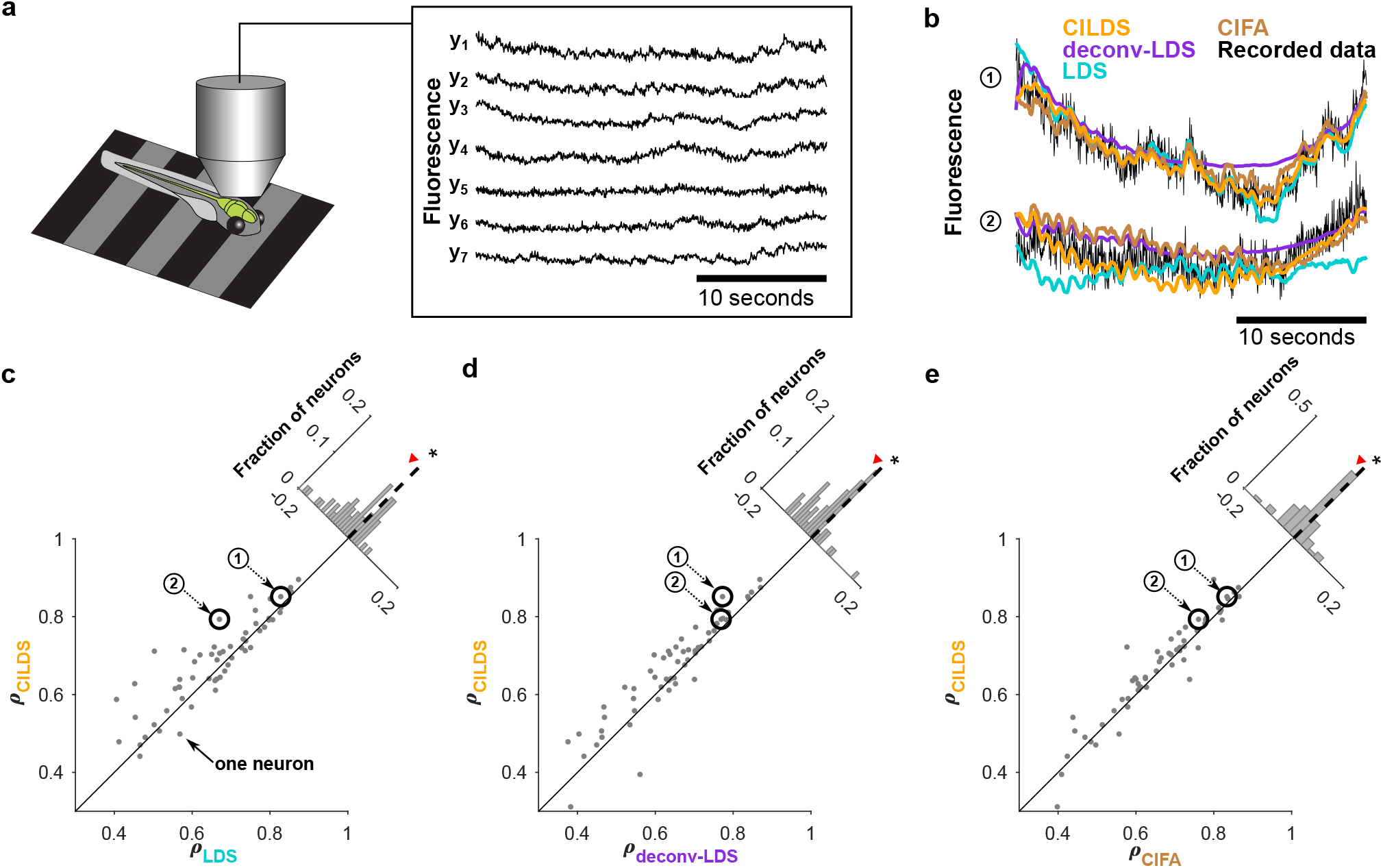
In larval zebrafish DRN recordings, CILDS captures the latent time courses better than the other three methods. **(a)** Two-photon calcium imaging using GCaMP6f at 30Hz was performed on three larval zebrafish in a virtual reality environment. Shown are representative fluorescence traces from seven of the imaged neurons. **(b)** Example recorded fluorescence traces (black) and leave-neuron-out predicted fluorescence using CILDS (orange), deconv-LDS (purple), LDS (cyan), and CIFA (brown). **(c-e)** Correlation between the recorded fluorescence and the leave-neuron-out predicted fluorescence for CILDS versus each of the other methods. The correlation is higher for CILDS than (c) LDS (*p <* 1 × 10^−7^, *n* =60 neurons, paired two-tailed t-test), (d) deconv-LDS (*p <* 1 × 10^−4^, *n* =60 neurons), and (e) CIFA (*p <* 0.05, *n* =60 neurons). Each point represents one neuron, where the correlation is computed for each trial (27 seconds long) then averaged across all 15 trials. The numbered points (black circles) correspond to the examples shown in panel (b). Diagonal histogram shows the paired difference in correlation between CILDS and one of the other methods, as indicated. A paired two-tailed t-test is applied to assess statistical significance, with statistical significance indicated by an asterisk. Note that the histogram is zoomed-in for visual clarity, and therefore the ends of the histograms are not shown.

The second experimental context involves awake, head-fixed mice passively viewing static visual gratings (Fig. 6a)^40^. Two-photon calcium imaging was performed on neurons expressing GCaMP6f in the primary visual cortex (V1) of three mice (133 to 319 neurons; mean: 234.7). Comparing the raw recorded fluorescence of the two experimental contexts (Fig. 5a versus Fig. 6a), the neurons in the DRN of the larval zebrafish tend to exhibit slower fluctuations in fluorescence that are correlated across neurons (Fig. 5a). In contrast, the neurons in mouse V1 exhibit faster changes in fluorescence, which appear less correlated across neurons (Fig. 6a). We applied the same analyses in Fig. 5 to these mouse recordings (Fig. 6b). Despite the stark differences between the fish DRN and mouse V1 fluorescence traces, we again found that CILDS more accurately predicted the fluorescence of held-out neurons than LDS (Fig. 6c; 63% of neurons above the diagonal), deconv-LDS (Fig. 6d; 68% of neurons above the diagonal), and CIFA (Fig. 6e; 82% of neurons above the diagonal). These results were also true for each mouse individually (Supplementary Fig. 6).

**Figure 6.**
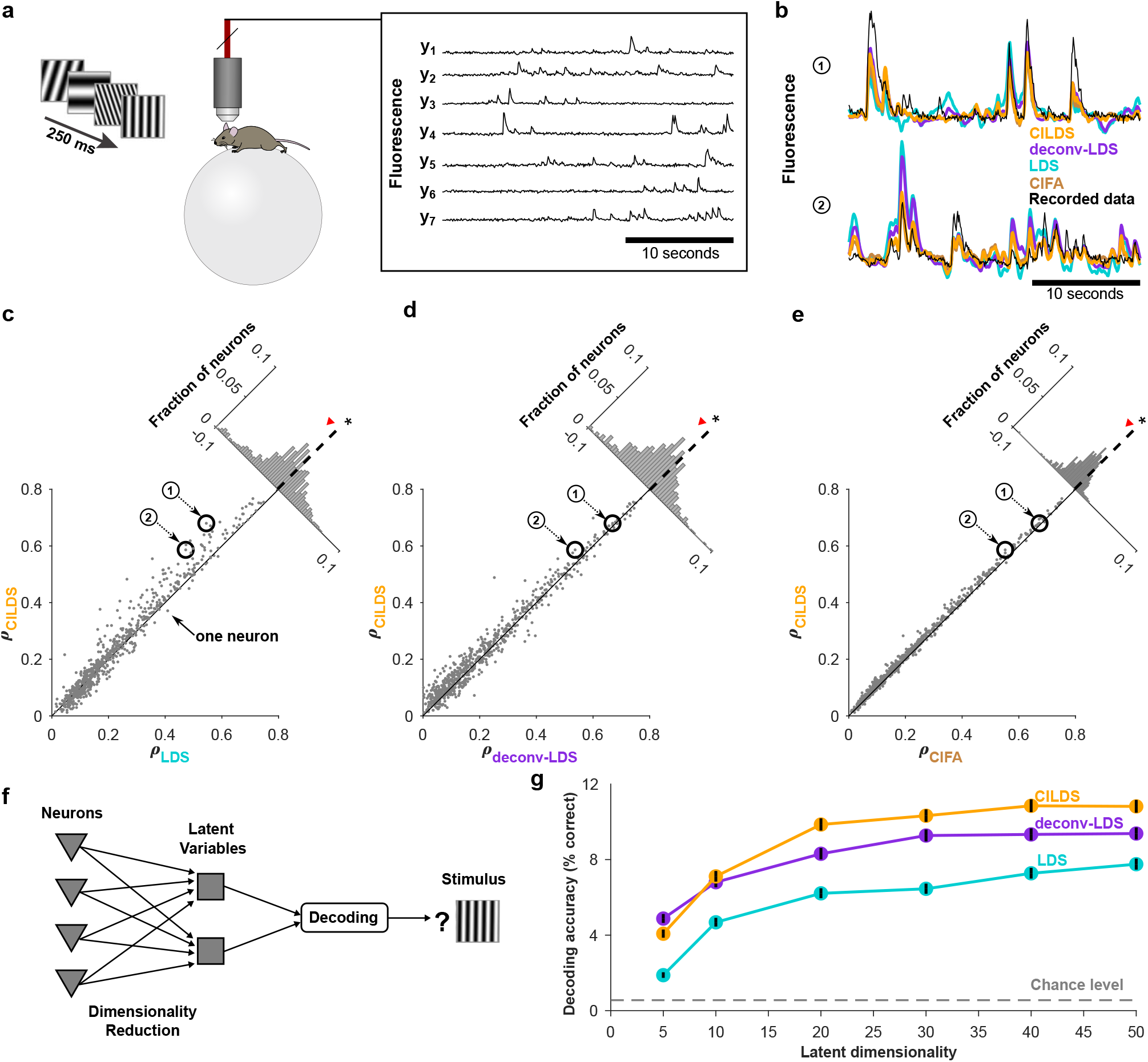
In mice V1 recordings, CILDS captures the latent time courses better than the other three methods. **(a)** Two-photon calcium imaging performed on awake mice viewing static gratings with different spatial frequencies and orientations (180 total stimuli) using GCaMP6f at 15.5Hz. Shown are representative fluorescence traces from seven of the imaged neurons. **(b)** Example segment of recorded fluorescence traces (black) and leave-neuron-out predicted fluorescence traces using CILDS (orange), deconv-LDS (purple), LDS (cyan), and CIFA (brown). **(c-e)** Correlation between recorded fluorescence and leave-neuron-out predicted fluorescence for CILDS versus each of the other methods. The correlation is higher for CILDS than (c) LDS (*p <* 1 × 10^−16^, *n* =704 neurons, paired two-tailed t-test), (d) deconv-LDS (*p <* 1 × 10^−26^, *n* =704 neurons), and (e) CIFA (*p <* 1 × 10^−80^, *n* =704 neurons). Each point represents one neuron, where the correlation is computed for each trial (196.7 seconds long) then averaged across all 15 trials. The numbered points (black circles) correspond to the examples shown in panel (b). Diagonal histogram shows the paired difference in correlation between CILDS and one of the other methods, as indicated. Asterisk denotes *p <* 0.05 for the paired two-tailed t-test above. Note that the histogram is zoomed-in for visual clarity, and therefore the ends of the histograms are not shown. **(f)** Flow diagram depicting decoding of visual stimuli using low-dimensional latent variables, which are obtained by applying a dimensionality reduction method to the recorded fluorescence traces. **(g)** Classification accuracy of the visual stimulus based on latent variables extracted using CILDS (orange), deconv-LDS (purple), and LDS (cyan). Classification was performed using a Gaussian Naive Bayes decoder, where the number of latent variables extracted by each dimensionality reduction method was systematically varied (horizontal axis). There were 180 total gratings (with different orientations and spatial frequencies) shown during the experiment, so the chance classification accuracy is 1/180 (gray dashed). The decoding window was 250 ms, which is the duration of stimulus presentation. Black error bars indicate indicate 95% confidence intervals (Bernoulli process).

Thus far, we have shown that CILDS extracts latent variables that more accurately predict the fluorescence of held-out neurons than the other methods. The leave-neuron-out evaluation assesses each method’s ability to capture the shared activity changes among neurons. Another way to assess how meaningful the latent variables extracted by each method are is to measure how strongly they reflect external variables, such as the sensory stimulus^41^. We thus performed a decoding analysis, whereby we classified the orientation and spatial frequency of the presented grating using the latent variables extracted by each of three methods (Fig. 6f). The stimuli could be more accurately decoded from the latent variables extracted by CILDS than from the latent variables extracted by either deconv-LDS or LDS (Fig. 6g). At the highest decoding accuracy of each method, CILDS outperformed deconv-LDS by 1.16 times and LDS by 1.43 times. This demonstrates that CILDS is better at capturing the shared activity changes among neurons that are relevant to the visual stimulus than the other two methods. Furthermore, the latent variables extracted by deconv-LDS provided more accurate decoding of the stimuli than the latent variables extracted by LDS, which indicates that accounting for the calcium transients is important when the variable of interest (here, the visual stimulus, which changes every 250 ms) changes on the timescale of tens to hundreds of milliseconds.

Taken together, the results based on calcium imaging recordings from two different recording regimes are consistent with what we identified in simulation. Namely, deconvolution should be used jointly with a dimensionality reduction method that incorporates a latent dynamical model (as in CILDS), particularly if the neural process of interest changes on a timescale of tens to hundreds of milliseconds.

## Discussion

In this work, we investigated the use of dimensionality reduction to summarize the time course of calcium imaging recordings. We considered dimensionality reduction approaches with and without deconvolution, and developed a novel method (CILDS) which jointly performs deconvolution and dimensionality reduction. We compared these methods using simulated fluorescence time courses, where we systematically varied experimental variables, as well as using calcium imaging recordings from larval zebrafish and mice. We found that: 1) CILDS, which leverages the population of neurons to jointly perform deconvolution and dimensionality reduction, outperformed the other methods, 2) using deconvolution was increasingly important with faster latent timescales, and 3) the use of a latent dynamical model, as in CILDS, was important for peering through the calcium decay transients to accurately recover the neural trajectories. Overall, this work provides a foundation for using dimensionality reduction to extract single-trial neural trajectories from calcium imaging recordings.

This work focused on the question of which dimensionality reduction method is most appropriate for extracting single-trial neural trajectories from calcium imaging recordings. There are two other settings in which dimensionality reduction is commonly used. First, dimensionality reduction can be applied to analyze how trial-averaged activity differs across experimental conditions (e.g., ref.^4,5,7,17^). In a recent study, Wei et al., applied principal components analysis (PCA) to trial-averaged electrophysiological recordings and calcium imaging with GCaMP6s^27^. They found important differences in the low-dimensional PCA trajectories obtained from electrophysiological recordings versus calcium imaging. This difference was mitigated by first deconvolving the calcium imaging recordings before applying dimensionality reduction, consistent with our findings. It may be possible to further improve the correspondence by applying CILDS to single-trial fluorescence recordings, then averaging the extracted low-dimensional neural trajectories across trials.

Second, dimensionality reduction is often used to analyze the trial-to-trial variability of neural population activity without time courses, i.e., using one time point or time window per trial (e.g., ref.^9,42–45^). In this case, there would be no information about how the calcium decays and so one would not be able to make use of a method that incorporates deconvolution. One might consider using CIFA, by analogy to the use of factor analysis to study the trial-to-trial variability of spike counts without time courses. However, it is important to note that, like CILDS, CIFA also requires multiple time points to be able to deconvolve, even though the latent variables in CIFA are independent from one time point to the next. If the original time series of calcium imaging is available, one can apply CILDS to the time series first, then average across the time points of the extracted latent variables. If the original time series of calcium imaging is not available, then a standard dimensionality reduction method such as factor analysis might be more suitable.

Previous studies have proposed methods for analyzing calcium imaging recordings that include latent variables and deconvolution in the same statistical model. Triplett et al.^46^ developed a method to study the interaction between evoked and spontaneous activity using calcium imaging recordings in sensory systems. In their model, the latent variables represent activity fluctuations shared amongst neurons that are not explained by the sensory stimulus, where these activity fluctuations are defined to be spontaneous activity. Aitchison et al.^47^ developed a method to infer spiking activity and neural connectivity from calcium imaging experiments that involve optogenetic stimulation. In their model, the latent variables represent shared activity amongst neurons that are not explained by the optogenetic stimulation or the activity of other neurons recorded simultaneously, and are intended to represent input from other brain areas. We developed CILDS for extracting latent variable time courses to summarize the population activity on individual experimental trials. In contrast to the two methods above which have more specific analysis goals (to separate evoked from spontaneous activity in the case of Triplett et al., or to infer spiking activity and neural connectivity in the case of Aitchison et al.), CILDS is general-purpose and well-suited for exploratory data analysis. This is akin to the use of methods such as LDS^48^, GPFA^37^, TCA^49^, LFADS^41^, and dPCA^50^ for exploratory analysis of population spiking activity to extract latent variable time courses, whose results can then lead to the use of population analysis methods with more specific goals (see ref.^51–53^ for examples).

CILDS can be extended in the following ways. First, the order of the autoregressive process for the calcium dynamics determines how quickly the calcium concentration (and consequently the fluorescence) rises after each spiking event^34^. Here, we used an autoregressive order of one, which corresponds to an instantaneous rise in calcium after each spiking event, as was done in previous work^32,34,54^. Although this is a reasonable first approximation, when imaging rates are fast or the calcium indicator is slow, it may be desirable to use an autoregressive order greater than one to better capture the non-instantaneous rise in calcium^13^. Second, different latent time series models can be used in the place of the linear dynamical system to achieve different analysis goals. For example, if one seeks only temporal smoothing in the latent variables without explicit dynamics, one can replace the linear dynamical system with Gaussian processes^37^.

Dimensionality reduction on population spiking activity has led to many insights about brain function. With the development of dimensionality methods that are tailored for calcium imaging, such as CILDS, we can leverage the advantages of calcium imaging such as being able to obtain information about neuron type or knowledge about which neurons project to other brain areas. For example, one could use dimensionality reduction to understand how populations of different neuron types interact^55^, or one could utilize information about where neurons project, coupled with dimensionality reduction, to understand how the projections contribute to the coordination of activity between brain areas^56,57^. This can enable novel insights about neural population activity recorded using calcium imaging that go beyond what is currently possible with electrophysiology.

## Methods

### Dimensionality reduction methods

Here we describe mathematically the dimensionality reduction methods used in this work: LDS, deconv-LDS, CILDS, and CIFA. For the purposes of this work, we assume that the spatial footprint of each neuron has already been identified from the raw calcium imaging data (a procedure known as image segmentation^54,58^), resulting in a fluorescence time course for each neuron. The dimensionality reduction methods presented here are applied to these fluorescence time courses.

#### Linear Dynamical System (LDS)

We first considered a standard dimensionality reduction method for summarizing the time course of spiking activity, a latent Linear Dynamical System (LDS), here applied to calcium imaging recordings. Let ***y***_*t*_ ∈ ℝ^*q*×1^ be a high-dimensional vector of fluorescence values recorded at time point *t*, where *q* is the number of neurons imaged simultaneously. The goal is to extract a corresponding low-dimensional latent variable ***z***_*t*_ ∈ ℝ^*p*×1^ at each time point, where *p* is the number of latent dimensions (*p < q*). The observation model is

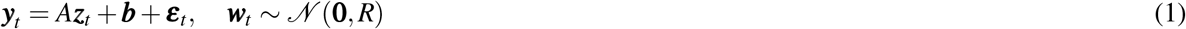

where *A* ∈ ℝ^*q*×*p*^ is the loading matrix that specifies how each neuron’s activity is related to the latent variables, ***b*** ∈ ℝ^*q*×1^ is an offset vector that accounts for constant background fluorescence, *R* ∈ ℝ^*q*×*q*^ is the observation noise covariance, and *t* = 1, …, *T*. We constrained *R* to be diagonal, thereby capturing activity variability and imaging noise independent to each neuron. The time-evolution of the latent variables is described as a linear dynamical system

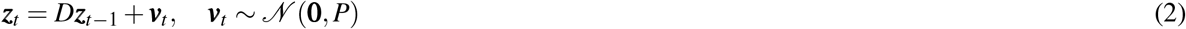

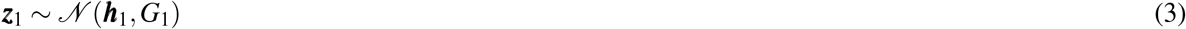

where *D* ∈ ℝ^*p*×*p*^ is the dynamics matrix that determines the timescale of the latent variables, *P* ∈ ℝ^*p*×*p*^ is the dynamics noise covariance, ***h***_1_ ∈ ℝ^*p*×1^ and *G*_1_ ∈ ℝ^*p*×*p*^ describe the mean and covariance of the latent variable at the first time point, and *t* = 2, …, *T*. We constrained *D, P*, and *G*_1_ here to be diagonal as a form of regularization, although a general LDS with these parameters unconstrained could also be used.

Equations (1), (2), and (3) together define the latent LDS^59^. We fit the model parameters (*A*, ***b***, *R, D, P*, ***h***_1_, *G*_1_) using the expectation-maximization (EM) algorithm^60^. To initialize the model parameters, we first performed factor analysis (FA) on ***y***_*t*_ to obtain *A*, ***b***, and *R*. We initialized *D* to be 0.999*I* (a stable system), which we found to work well in practice in the simulations. We ran the EM algorithm until convergence (defined as a log data likelihood increase of *<* 10^−6^ or 1500 iterations, whichever came first). This maximum number of iterations was chosen heuristically, by noting empirically that the latent variables do not change significantly beyond this point.

#### Deconvolution - Linear Dynamical System (deconv-LDS)

Since fluorescence traces are an indirect measure of spiking activity, we also considered a two-stage approach, whereby we first deconvolve the fluorescence traces one neuron at a time to estimate the underlying spiking activity. Then we applied a standard dimensionality reduction method, in this case LDS, to those deconvolved estimates. We refer to this two-stage method as deconv-LDS.

For the deconvolution stage of deconv-LDS, we used the Online Active Set method to Infer Spikes (OASIS), developed by Friedrich et al.^34^, using their *L*_1_ regularization. We also tested their *L*_0_ regularization and found the *L*_1_ regularization to work better for our datasets. As per Friedrich et al., this first order autoregressive model for OASIS is described for each neuron as

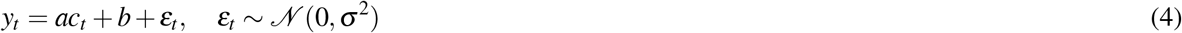

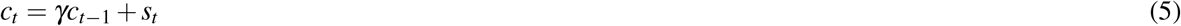

where *y*_*t*_ is the recorded fluorescence at at time *t, c*_*t*_ represents the calcium concentration at time *t, ε*_*t*_ captures imaging noise independent of the calcium and spiking activity, and *s*_*t*_ is the spiking activity. The parameter *a* relates the calcium concentration to fluorescence, *b* accounts for the baseline fluorescence, *σ*^2^ captures the variance of the imaging noise, and *γ* specifies how quickly the calcium trace decays, which depends on the calcium indicator. Additionally, there is a hyperparameter in the OASIS model, minimum spike size, that sets the minimum value of *s*_*t*_ that would be identified. In this model, all variables are scalars and *a* is constrained to be non-negative. We initialized OASIS with *γ* values that are typical for the calcium indicators used^13^ (see Supplementary Table 1), and allowed OASIS to optimize *a, b, γ, σ*^2^ and the minimum spike size.

Applying deconvolution to the recorded fluorescence traces returned estimates of the time course of spiking activity for each neuron, *s*_*t*_. We then used the estimated spiking activity of all the neurons as the observations ***y***_*t*_ ∈ ℝ^*q*×1^ for time points *t* = 1, …, *T* in the LDS model defined in equations (1)-(3).

#### Calcium Imaging Linear Dynamical System (CILDS)

CILDS unifies the approaches described above by allowing estimates of shared activity among neurons (i.e., the latent variables) to influence the estimates of deconvolved spiking activity, and vice versa. In other words CILDS performs deconvolution for all neurons and dimensionality reduction jointly, in a unified framework. This is in contrast to deconv-LDS, which deconvolves the activity of each neuron independently. With low-dimensional latent variables that are jointly estimated with the model of calcium decay, CILDS is better able to peer through the calcium decay to more clearly identify the shared activity among neurons, as compared to deconv-LDS and LDS applied directly on fluorescence.

Let ***y***_*t*_ ∈ ℝ^*q*×1^ be the high-dimensional vector of fluorescence traces recorded at each time point *t*, where *q* is the number of neurons imaged simultaneously. The goal is to extract a corresponding low-dimensional latent variable ***z***_*t*_ ∈ ℝ^*p*×1^ at each time point, where *p* is the number of latent dimensions (*p < q*). The observation model follows the multivariate form of equation (4) which was used for deconvolution

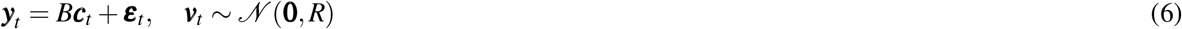

where *B* ∈ ℝ^*q*×*q*^ maps the calcium concentration to the recorded fluorescence, *R* ∈ ℝ^*q*×*q*^ is the fluorescence noise covariance, and *t* = 1, …, *T*. We constrained *B* and *R* to be diagonal to allow each dimension of ***c***_*t*_ to represent the calcium concentration of one neuron. *B* accounts for all experimental variables influencing the scale of the signal from each neuron, such as the amplification of the imaging system^32^, and *R* accounts for fluorescence fluctuations independent of calcium concentration. Here we omit the additive offset found in equation (4) without loss of generality due to the offset included in equation (7).

Similar to equation (5), the calcium decay for each neuron is described using a first order autoregressive model

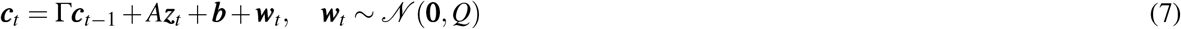

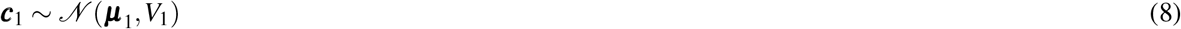

where Γ ∈ ℝ^*q*×*q*^ captures the calcium decay, *A* ∈ ℝ^*q*×*p*^ is the loading matrix that describes how latent variable maps to calcium concentrations, ***b*** ∈ ℝ^*q*×1^ is a constant vector, *Q* ∈ ℝ^*q*×*q*^ captures the spiking variability independent to each neuron, ***µ***_1_ ∈ ℝ^*q*×1^ and *V*_1_ ∈ ℝ^*q*×*q*^ describe the mean and variance of the calcium concentration at the first time point, respectively, and *t* = 2, …, *T*. Our key innovation is to replace the spiking activity *s*_*t*_ from equation (5) with *A****z***_*t*_ + ***b*** + ***w***_*t*_. By analogy to factor analysis, *A****z***_*t*_ describes the shared activity among neurons, and ***w***_*t*_ describes the activity independent to each neuron. We constrain Γ, *Q*, and *V*_1_ to be diagonal to prevent intermixing among neurons outside of the loading matrix. The form of Γ allows each neuron to have a different calcium decay as determined by the fitting procedure (see below), which can depend on the extent of calcium buffering within a cell and the calcium indicator used^13,32,34^.

Similar to LDS, the low-dimensional latent variables ***z***_*t*_ evolve over time according to a linear dynamical system

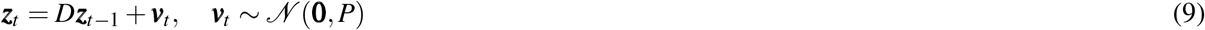

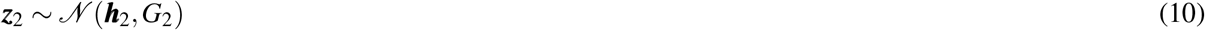

where *D* ∈ ℝ^*p*×*p*^ is the dynamics matrix, *P* ∈ ℝ^*p*×*p*^ is the dynamics noise covariance, ***h***_2_ ∈ ℝ^*p*×1^ and *G*_2_ ∈ ℝ^*p*×*p*^ are the mean and covariance of the latent variable at the first time point respectively, and *t* = 3, …, *T*. We constrained *D, P*, and *G*_2_ here to be diagonal as a form of regularization, although a model with these parameters unconstrained could also be used. Note that according to equation (7), ***z***_2_ is the first latent variable in the time series (i.e., there is no ***z***_1_).

Equations (6) - (10) define CILDS. The joint estimation of the parameters *B, R*, Γ, *A*, ***b***, *Q*, ***µ***_1_,*V*_1_, *D, P*, ***h***_2_, and *G*_2_ allows CILDS to leverage the entire recorded neural population to perform deconvolution and estimate latent variables in a unified fashion. CILDS can be viewed as a special case of the standard LDS, where the parameters are constrained in a specific way. We fit CILDS using the EM algorithm, initialized using deconv-LDS run for 100 EM iterations. The EM algorithm was run until convergence, defined as a log data likelihood increase of *<* 10^−6^ or 1500 iterations, whichever came first. The maximum number of iterations was chosen by noting empirically that the latent variables do not change substantially beyond this point. The EM equations for CILDS are provided in Supplemental Information.

#### Calcium Imaging Factor Analysis (CIFA)

To evaluate the importance of incorporating a latent dynamical system in dimensionality reduction methods for calcium imaging, we developed a method (CIFA) identical to CILDS, with the only difference being that CIFA does not enforce latent dynamics. Like CILDS, CIFA also uses equations (6), (7), and (8). The only difference between CIFA and CILDS is that we replaced the latent dynamical system equations (9) and (10) with

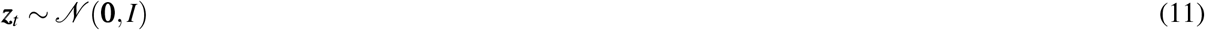

for *t* = 2, …, *T*. In other words, CIFA defines latent variables that are independent across time, whereas CILDS defines latent variables that evolve smoothly over time. Like CILDS, we fit CIFA using the EM algorithm, initialized using parameters from deconv-FA run for 100 EM iterations. The EM algorithm was run until convergence, defined as a log data likelihood increase of < 10^−6^ or 1500 iterations, whichever came first. The EM equations for CIFA are the same as for CILDS (see Supplemental Information), without the equations estimating *D, P*, ***h***_2_, *G*_2_. Note that there is no loss of generality by setting the prior distribution of ***z***_*t*_ to 𝒩 (**0**, *I*), compared to a general Gaussian distribution. Additionally, although CIFA has FA in its name, there is one key difference from FA. Whereas FA can be fit on data with no concept of time, CIFA requires a time series for deconvolution (equations (7) and (8)).

### Simulation framework

We created a framework to simulate fluorescence recordings from calcium imaging for two reasons. First, in calcium imaging recordings, the ground truth latent variables are unknown. A simulation of fluorescence traces from known latent variables allows us to directly evaluate our dimensionality reduction methods by comparing the estimated latent variables with the ground truth latent variables. Second, we wanted a simulation framework in which we could systematically vary various experimentally relevant parameters to see their effects on the estimated latent variables. Specifically, we evaluated our ability to recover the ground truth latent variables as a function of the timescale of the latent variables, the number of neurons, the calcium decay rate, and the size of the fluorescence noise. The simulation procedure consists of first generating fluorescence traces from known latent variables while varying the experimentally relevant parameters listed above. Then, we applied each dimensionality reduction method to estimate latent variables from the simulated fluorescence traces. The estimated latent variables were then compared to the ground truth latent variables. The steps of this procedure are detailed below.

#### Generating fluorescence traces

To simulate fluorescence traces, we first drew *p* Gaussian processes (GP)^61^, where each GP has *T* time points at 1 ms time resolution. We denote the *i*th GP as ***z***_*i*,:_ ∈ ℝ^1×*T*^, where *i* = 1, …, *p*. The GP allows us to specify the covariance *K*_*i*_ ∈ ℝ^*T* ×*T*^ for the *i*th GP across the *T* time points as

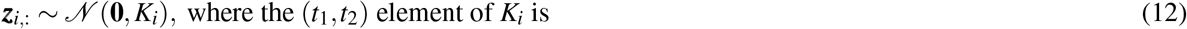

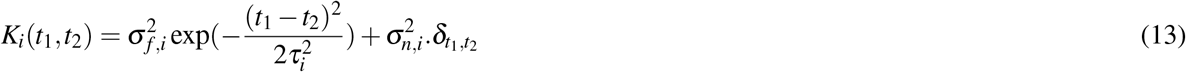

and *t*_1_,*t*_2_ = 1, …, *T*. Here we chose the commonly used squared exponential covariance function. The squared exponential covariance is defined by its signal variance 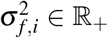, characteristic timescale *τ*_*i*_ ∈ ℝ_+_, and noise variance 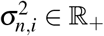. The Kroneker delta 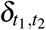 equals 1 if *t*_1_ = *t*_2_ and 0 otherwise. We set 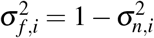 so that thelatent variable ***z***_*t*_ ∈ ℝ^*p*×1^at every time point has mean **0** and a variance *I*. The vector ***z***_*t*_ comprises the *t*th time point from each of the *p* Gaussian processes. The noise variance 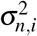 must be nonzero to ensure that *K*_*i*_ is invertible, hence for practical purposes we set 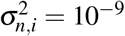. We note that an LDS can also be used to introduce latent dynamics, but we chose to use a GP due to the ease by which we can generate stationary time series with specified time scales. Furthermore, using a GP introduced model mismatch for all of the dimensionality reduction methods, which enables a more meaningful comparison across methods.

Next, we projected the low-dimensional latent variables ***z***_*t*_ into the high-dimensional neural space to obtain neural firing rates for each of *q* neurons at each time point *t* = 1, …, *T*. We then imposed a rectifying nonlinearity applied element-by-element using log(**1** + exp(*W* ***z***_*t*_ + ***µ***)), where *W* ∈ ℝ^*q*×*p*^ is the loading matrix and ***µ*** ∈ ℝ^*q*×1^ is a constant offset, to ensure that firing rates are non-negative. We generated binary spikes ***s***_*t*_ ∈ ℝ^*q*×1^ at each time point using an inhomogeneous Poisson process with time-varying rates defined by the output of this rectifying nonlinearity. Finally, we obtained the calcium concentration ***c***_*t*_ ∈ ℝ^*q*×1^ using a first order autoregressive model and fluorescence ***y***_*t*_ ∈ ℝ^*q*×1^, as in Friedrich et al. 2017^34^

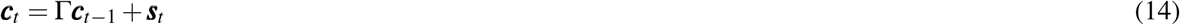

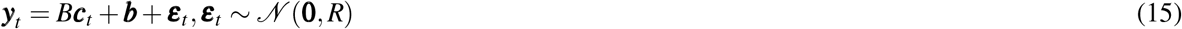

where Γ ∈ ℝ^*q*×*q*^ determines how quickly the calcium decays, *B* ∈ ℝ^*q*×*q*^ relates the calcium concentration to the fluorescence, ***b*** ∈ ℝ^*q*×1^ is the baseline fluorescence, *R* ∈ ℝ^*q*×*q*^ is the imaging noise covariance, and *t* = 1, …, *T*, at the same (1ms) resolution as the GP. We set Γ, *B*, and *R* to be diagonal. We specified Γ such that the decay constants approximately matched the decay constants of GCaMP6f, GCaMP6m, and GCaMP6s, found in Chen et al.^13^ (See Supplementary Table 1). *B* represents experimental variables influencing the scale of the calcium signal of each neuron, such as the amplification of the imaging system. *B* is set as the identity matrix and ***b*** is set to be **0** in our simulations. We varied the signal-to-noise ratio by varying *R* to simulate low, medium, and high noise regimes (Supplementary Table 1).

For each simulation run, we simulated *p* = 10 latent variables, where each latent dimension had the same latent timescale for ease of interpretation. Across different settings of experimentally relevant parameters, we explored a range of timescales *τ* ∈ {50,100,200,1000,2000,5000} ms. To define loading matrices *W* that were realistic for spiking activity, we used electrophysiological recordings with 94 neurons^6^. Across runs, we tested *q* ∈ {20,50,94} neurons. For the *q* = 20 and *q* = 50 cases, we randomly subsampled the electrophysiological recordings. To avoid a start-up transient at the start of every trial from the calcium model, we generated two long fluorescence traces for each neuron, each 6,000,000 time points long (1 ms resolution). One fluorescence trace was used for training, and the other for testing. We divided each trace into 100 trials. Each trial was 60 s long, comprising 60,000 time points. Finally, having generated these fluorescence traces at 1 kHz, we down-sampled the fluorescence rate to a more typical imaging rate of 40 Hz (2,400 time points per trial). The simulation parameters are summarized in Supplementary Table 1. See Practical Considerations below for how we generated GPs with 6,000,000 time points within the computer’s memory constraints.

#### Evaluation of dimensionality reduction methods

We applied the four dimensionality reduction methods to the simulated fluorescence traces to evaluate how accurately each method reconstructed the ground truth latent variables. We used two-fold cross validation in these estimates, so that the model parameters were fit using half of the data (100 training trials, each with 2,400 time points) and those parameters were then used to estimate the latent variables in the other half of the data (100 held-out trials). For all methods, the latent variables are only unique up to an arbitrary linear transformation. In order to compare the estimated and ground truth latent variables, we aligned them using the following procedure. First we split the estimated latent variables for each test fold into two further inner halves. We concatenated the trials over time, such that the estimated latent variables are defined as 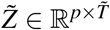, and the ground truth latent variables are defined as 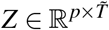, where *p* = 10 is the dimensionality of the latent variables and 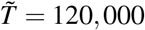 for one inner half. Then we applied linear regression to relate 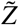 and the *Z* corresponding to that inner half. This yielded a transformation matrix *W*, where 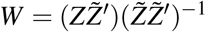. Finally we used *W* to align the 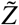 corresponding to the other inner half to obtain the transformed estimated latent variables 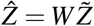, where 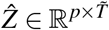. We performed the same procedure to align each inner half of each cross-validation fold.

We computed the accuracy of the *i*th transformed estimated latent variables

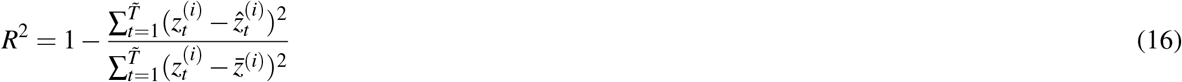

where 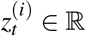 is the (*i,t*) entry of *Z*, and 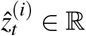 is the (*i,t*) entry of 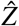. The mean of the *i*th ground truth latent variable across time is defined as 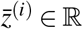, where *i* = 1, …, *p*. A larger *R*^2^ means a better match between the estimated latent variables and the ground truth latent variables, based on the proportion of total variance (of the ground truth latent variables) explained. *R*^2^ has an upper limit of 1, and any value below 0 indicates that the estimate is poorer than the using the ground truth mean. We repeated this process for each cross-validation fold and averaged the results across all *p* latent dimensions and folds. It is important to note that since the results are cross-validated, a dimensionality reduction method with more parameters will not necessarily outperform a method with fewer parameters.

### Experimental data

#### Larval Zebrafish

Neurons were imaged from the dorsal raphe nucleus (DRN) of larval zebrafish while they engaged in a “fictive swimming” motosensory gain adaptation task^39^. Calcium imaging was performed using light-sheet microscopy at 30 Hz on a single plane of narrow area around the DRN. This was performed with *Tg(elavl3:GCaMP6f)*^*jf1*^ fish expressing GCaMP6f in the cytosol^62^. We analyzed recordings from three fish. In the task, the fish underwent an initialization period of 20 seconds to increase locomotor drive, a training period in which the fish attenuated their locomotor drive, and a delay period of 10 seconds which stopped the fish from swimming. Finally, there was a test period of 5 seconds to probe the extent to which the attenuated locomotor drive persisted throughout the delay period. There were three different training period lengths of 7, 15, or 30 seconds. Here we combined the data from the different training periods. We included only the 20-second initialization period and the first 7 seconds of the training period of each trial. Thus, the analyzed portion of each trial is nominally identical. For each fish, this yielded 15 trials, each 27 seconds long. We analysed 22, 19, and 19 neurons imaged from the DRN of each fish, respectively.

#### Mouse

Two-photon calcium imaging was performed in the binocular zone of V1 in awake head-fixed mice resting atop a floating spherical treadmill^40^. GCaMP6f was expressed in excitatory neurons and imaged at 15.5 Hz. We analyzed recordings from three mice. Mice were positioned to passively view static sinusoidal gratings, without reward. There were 180 gratings presented, comprising 12 different orientations equally spaced with range {0 − 165}° and 15 different spatial frequencies equally spaced with range {0.02 − 0.30} cycles/°. Each “trial” was a 196.7 seconds long recording (3049 time points), comprising 4 presentations of each of the 180 possible gratings in random order. Each presentation lasted 250 ms without an intervening grey screen. The onset time of the first stimulus relative to the beginning of the recording was varied. This means that there is a short period of time recorded before the first stimulus is shown, and a short period of time recorded after the last stimulus is shown. The experiment comprised 15 trials for each mouse. We analysed 133, 252, and 319 neurons from V1 of mouse (labeled mouse 1-3, respectively). These mice correspond to mouse2317, mouse2320, and mouse2209 in the experiments.

### Data analysis

#### Leave-neuron-out fluorescence prediction

We sought to compare the four dimensionality reduction methods using experimental data. In experimental data, ground truth latent variables are unknown, and so we could not use the same evaluation procedure of comparing estimated and ground truth latent variables as in the simulations. Furthermore, the cross-validated data likelihoods are not comparable across all methods. Hence to compare the four methods, we performed a leave-neuron-out fluorescence prediction test to determine which method best summarizes the neuronal activity with low-dimensional latent variables^37,38^. The intuition is that a method that provides a better summary of the population activity using the latent variables would be better able to predict the activity of held-out neuronal fluorescence traces.

For the leave-neuron-out fluorescence prediction, we performed 5-fold cross-validation. We first split the trials into five equal-sized folds. For a given dimensionality reduction method, we fit the model parameters using four of the folds. With the remaining validation fold, we estimated the latent variables using all but one neuron, and then predicted the activity of that held-out neuron using those estimated latent variables. We did this for every neuron in the validation fold. In the same manner, we performed leave-neuron-out fluorescence predictions for the remaining folds and thus obtained predictions of the fluorescence activity for all the neurons at all time points. We then computed the Pearson’s correlation coefficient between the predicted and recorded fluorescence traces.

Performing the leave-neuron-out prediction procedure requires selecting the latent dimensionality *p* for each method. For the larval zebrafish DRN recordings, we used nested cross-validation to select the optimal dimensionality for each method. The leave-neuron-out prediction then used this optimal latent dimensionality, which can be different for each method. Nested cross-validation uses the 5-fold cross-validation described earlier as the outer folds. Within each outer fold, the training portions are then used to perform an inner 4-fold cross-validation where we determined the optimal latent dimensionality using cross-validated data likelihood. We used this optimal latent dimensionality to estimate the model parameters using the same four training folds. These model parameters were then used for the leave-neuron-out prediction procedure of the remaining (validation) outer fold. Note that although the cross-validated data likelihood is not comparable across methods, it is comparable across different latent dimensionalities for the same method.

For the mouse V1 recordings, we found that the performance of all dimensionality reduction methods increased as latent dimensionality increased within the range of dimensionalities tested (5-50). Thus we used a latent dimensionality of 50 for all methods in our 5-fold cross validation procedure for leave-neuron-out prediction.

#### Decoding analysis

Another way to assess how meaningful the extracted latent variables are is by decoding external variables from the latent variables^41^. For the larval zebrafish recordings, there is no moment by moment behavior that can be decoded from the neural activity. In the mouse recordings, we can decode the orientation and spatial frequency of the grating stimuli. To begin, we applied the dimensionality reduction methods to the mouse recordings. We then applied a linear Gaussian Naïve Bayes classifier^63^ to the extracted latent variables

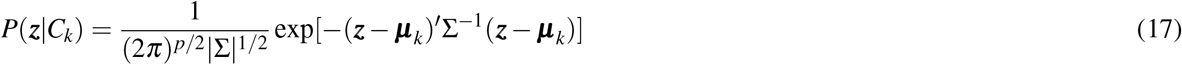

where ***z*** ∈ ℝ^*p*×1^ is the latent variable averaged across the time points in a 250 ms window corresponding to a given stimulus *C*_*k*_, *k* = 1, …, 180. The parameters of the classifier are ***µ***_*k*_, the mean of the latent variables corresponding to class *k*, and Σ, the covariance of the latent variables across trials. We constrain Σ to be diagonal and the same across all classes. The parameters ***µ***_*k*_ and Σ are fit by maximizing the likelihood of the training data using 5-fold cross-validation.

We then used the parameters found from the training data to classify the held-out data according to

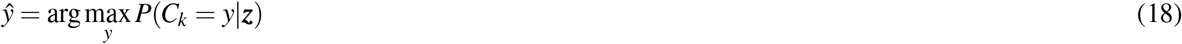

where ŷ is the predicted class label (1,…,180). We then computed the accuracy of the predicted class labels against the true class labels (chance level is 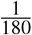). A higher accuracy indicates that the latent variables are better at capturing the shared modulations among neurons that are relevant to the visual stimulus. For latent dimensionalities {5,10,20,30,40,50}, we fit each dimensionality reduction method using all data. Then, we performed 5-fold cross-validation in the decoding stage.

To decode visual stimuli, there are two considerations. First, visual information takes time to arrive in the visual cortex, hence there is a need to shift the window of neural activity relative to the stimulus presentation. Second, there is an additional time delay introduced by the calcium indicators. Unlike deconv-LDS and CILDS, LDS does not attempt to remove the calcium decay. Hence, we expected a longer latency for LDS than for deconv-LDS and CILDS. To determine the appropriate window, we considered a range of time lags and evaluated their cross-validated classification accuracy. We found that the best cross-validated accuracy was obtained for a 4 time point (260 ms) shift for LDS, and a 3 time point (190 ms) shift for deconv-LDS and CILDS. Thus we used these time lags to report the classification accuracy of our decoding analysis (Fig. 6g).

### Practical considerations

#### Approximating a long Gaussian process

To carry out computations using a GP, one would need to represent a covariance matrix of size *T* × *T* in memory, where *T* =6,000,000 in our simulations. For such values of *T*, the memory requirement can exceed the memory capacity of the computer. To overcome this, we employed two strategies in tandem. First, we generated the GPs one segment at a time, rather than all *T* time points at once. For example, to generate a GP with 6,000,000 time points, we can first generate a GP with 5,000 time points according to equations (12) and (13). Then, we can generate a GP for the next 5,000 time points conditioned on the first 5,000 time points using Gaussian conditioning^61^

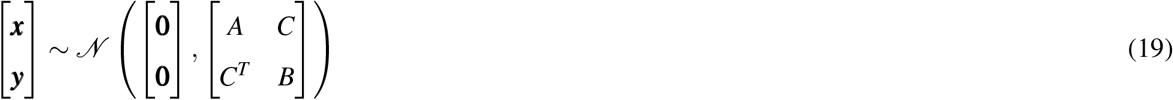

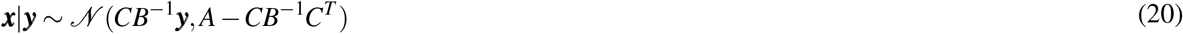

where *A* ∈ ℝ^5000×5000^ and *B* ∈ ℝ^5000×5000^ are the covariances of the second and first half of the GP, respectively. *C* ∈ ℝ^5000×5000^ is the covariance between the first and second half of the GP. We can continue this procedure, where for each new segment we condition on all of the segments that have been generated thus far. Statistically this procedure is equivalent to if we had generated the entire GP time series at once.

As we continue this procedure, the number of time points being conditioned on will grow and the matrices *B* and *C* in equations (19) and (20) can exceed the memory capacity of the computer. We thus employ a second strategy that leverages the fact that, according to the squared exponential covariance (13), two time points covary highly when they are close in time and almost independent when they are far apart in time. Thus, we make the approximation that in equations (19) and (20), we condition only on the most recent 5,000 time points.

#### Computational running time

The following are representative running times to fit the different dimensionality reduction models. These running times are based on single threads run on Matlab (2019a) using Intel(R) Xeon(R) CPU processors (Gold 6230, 2.1 GHz) with 250 GB of RAM. First, consider one cross-validation fold of the zebrafish recordings with 4 trials, 22 neurons, 10 latent variables, and 1950 time points per trial. For each EM iteration, CILDS takes on average approximately 0.9 s, LDS (as well as the second-stage of deconv-LDS) takes 0.4 s. Second, consider one cross-validation fold of the mouse recordings with 12 trials, 319 neurons, 30 latent variables, and 3049 time points per trial. For each EM iteration, CILDS takes on average approximately 110 s, and LDS (as well as the second-stage of deconv-LDS) takes 10 s. For all methods, the most expensive computations are the matrix inversions in the expectation step of the EM algorithm. There is a *p* × *p* matrix inversion at each time point for LDS (and deconv-LDS), and a (*p* + *q*) × (*p* + *q*) matrix inversion at each time point for CILDS, where *p* is the latent dimensionality and *q* is the number of neurons.

## Data Availability

The data that support the findings of this study are available from the authors upon reasonable request.

## Code Availability

Matlab code for the simulations and dimensionality reduction methods in this work will be made publicly available upon publication.

## Acknowledgements

The authors would like to thank Katrina P. Nguyen for the animal illustrations. This work was supported by the Agency for Science, Technology and Research (A*STAR) Singapore (T. Koh), Howard Hughes Medical Institute (W.E.B., T. Kawashima, and M.B.A.), Simons Foundation Simons Collaboration on the Global Brain Award 542943 (M.B.A.) and 543065 (B.M.Y.), the Shurl and Kay Curci Foundation (S.M.C. and S.J.K.), NIH R01 HD071686 (S.M.C. and B.M.Y.), NIH R01EY024678 (S.J.K.), NSF NCS BCS1533672 (S.M.C. and B.M.Y.), NSF CAREER award IOS1553252 (S.M.C.), NSF NCS BCS1734916 (B.M.Y.), NIH CRCNS R01 NS105318 (B.M.Y.), NIH CRCNS R01 MH118929 (B.M.Y.), NIH R01 EB026953 (B.M.Y.).

## Author contributions statement

T. Koh, W.E.B., S.M.C., and B.M.Y. designed the dimensionality reduction and analysis methods. T. Koh derived and implemented the dimensionality reduction methods. T. Koh performed the analyses, based on earlier analyses by R.S. T. Kawashima and M.B.A. contributed to evaluation of the methods. T. Kawashima, B.B.J., S.J.K., M.B.A. designed the animal experiments. T. Kawashima performed the larval zebrafish experiments, and B.B.J. performed the mouse experiments. T. Koh, W.E.B., M.B.A., S.M.C., and B.M.Y. wrote the manuscript. All authors discussed the results and commented on the manuscript. S.M.C. and B.M.Y. contributed equally to this work.

## Competing interests

The authors declare no competing interests.

## Supplementary Figures

**Supplementary Figure 1.**
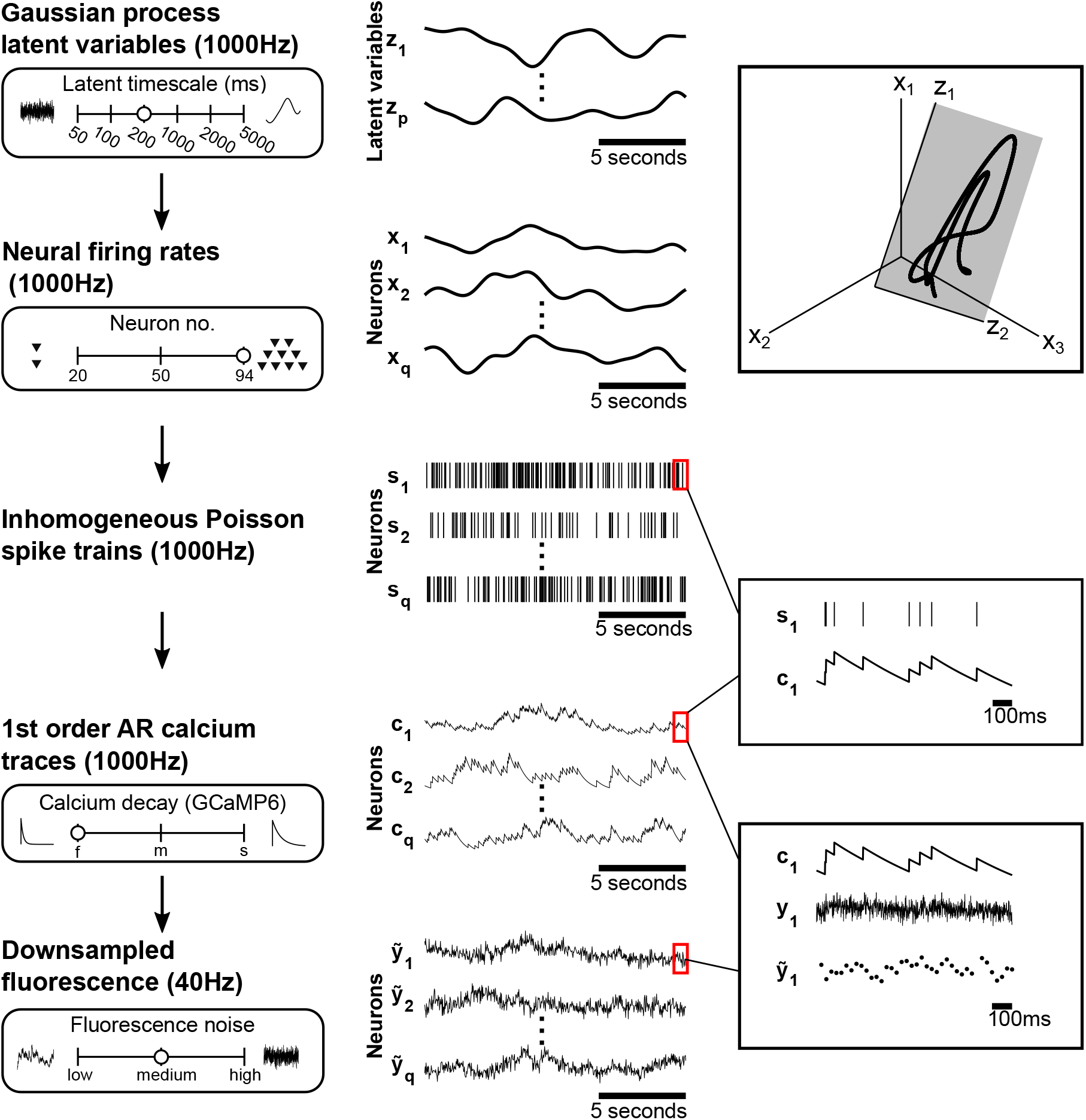
Simulation framework and range of parameters. Detailed simulation framework depicting the generation of fluorescence traces from latent variables. The sliders show the range of parameters explored, and the open circles represent one example of a set parameter values. The middle column shows example signals generated at each stage. Gaussian process latent variables ***z*** ∈ ℝ^*p*×*T*^ (sampled at 1000 Hz time resolution) are projected to form firing rates ***x*** ∈ ℝ^*q*×*T*^, where *p* is the number of latent variables, *q* is the number of neurons, and *T* is the number of time points in each trace. The upper right panel shows an example of the latent space (***z***) placed within a higher-dimensional neural space (***x***). Firing rates are used to generate spike trains, ***s*** ∈ ℝ^*q*×*T*^, according to an inhomogeneous Poisson process. Calcium traces, ***c*** ∈ ℝ^*q*×*T*^, are generated from spike trains using a 1st order autoregressive process^32^, with the center right panel showing a close-up of the spike-to-calcium transformation. White noise is added to calcium traces to form fluorescence traces ***y*** ∈ ℝ^*q*×*T*^. These traces are down-sampled to 40Hz, forming 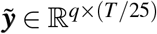, to more closely match typical sampling rates in calcium imaging recordings. The lower right panel shows a zoomed in view of this process.

**Supplementary Figure 2.**
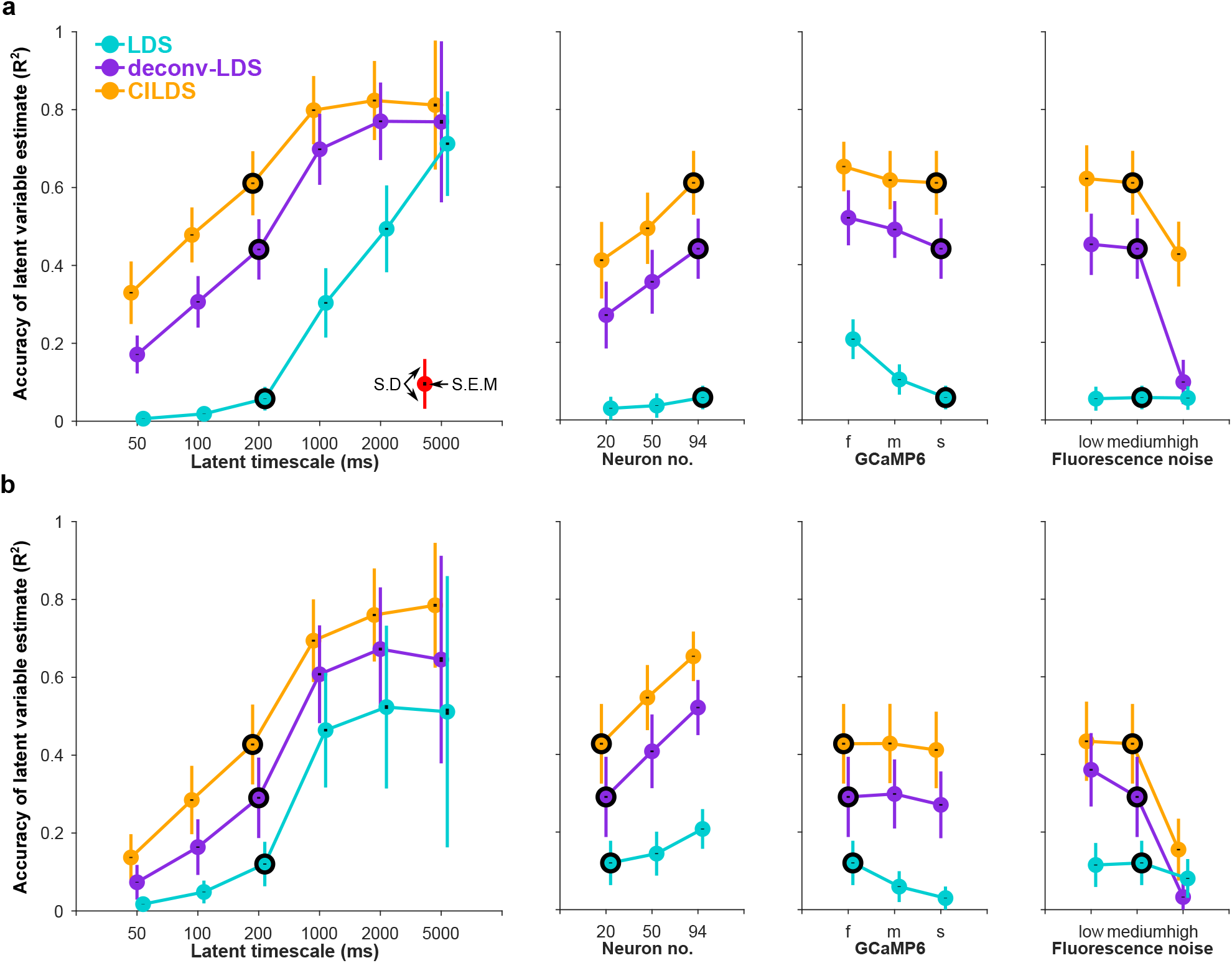
CILDS continues to outperform other methods under different settings of the experimental variables. Same conventions as Fig. 3d-g. These experimental variables are different from Fig. 3 in the following ways: **(a)** a calcium decay constant matching the calcium indicator GCaMP6s instead of GCaMP6f and **(b)** a smaller number of neurons (20 instead of 94). Overall CILDS extracts latent variables that more closely match the ground truth simulated latent variables than the other two methods, consistent with Fig. 3. Black circles represent the parameter settings that are fixed across all panels in each row. Comparing to Fig. 3, there are two notable features. First, when the calcium indicator decay is slow, it becomes even more important to deconvolve. Recall that CILDS and deconv-LDS both include deconvolution, whereas LDS does not. CILDS performs similarly whether the calcium indicator is fast (Fig. 3d) or slow (here in panel a). The same is true for deconv-LDS. By contrast, LDS performs worse for slow compared to fast calcium indicator decay because it does not include deconvolution. Second with fewer neurons, the performance of all methods goes down. And as a result, there is a smaller difference in performance between methods (here in panel b). With less statistical power to leverage for separating calcium decay and latent timescale, the three methods show less distinction in performance.

**Supplementary Figure 3.**
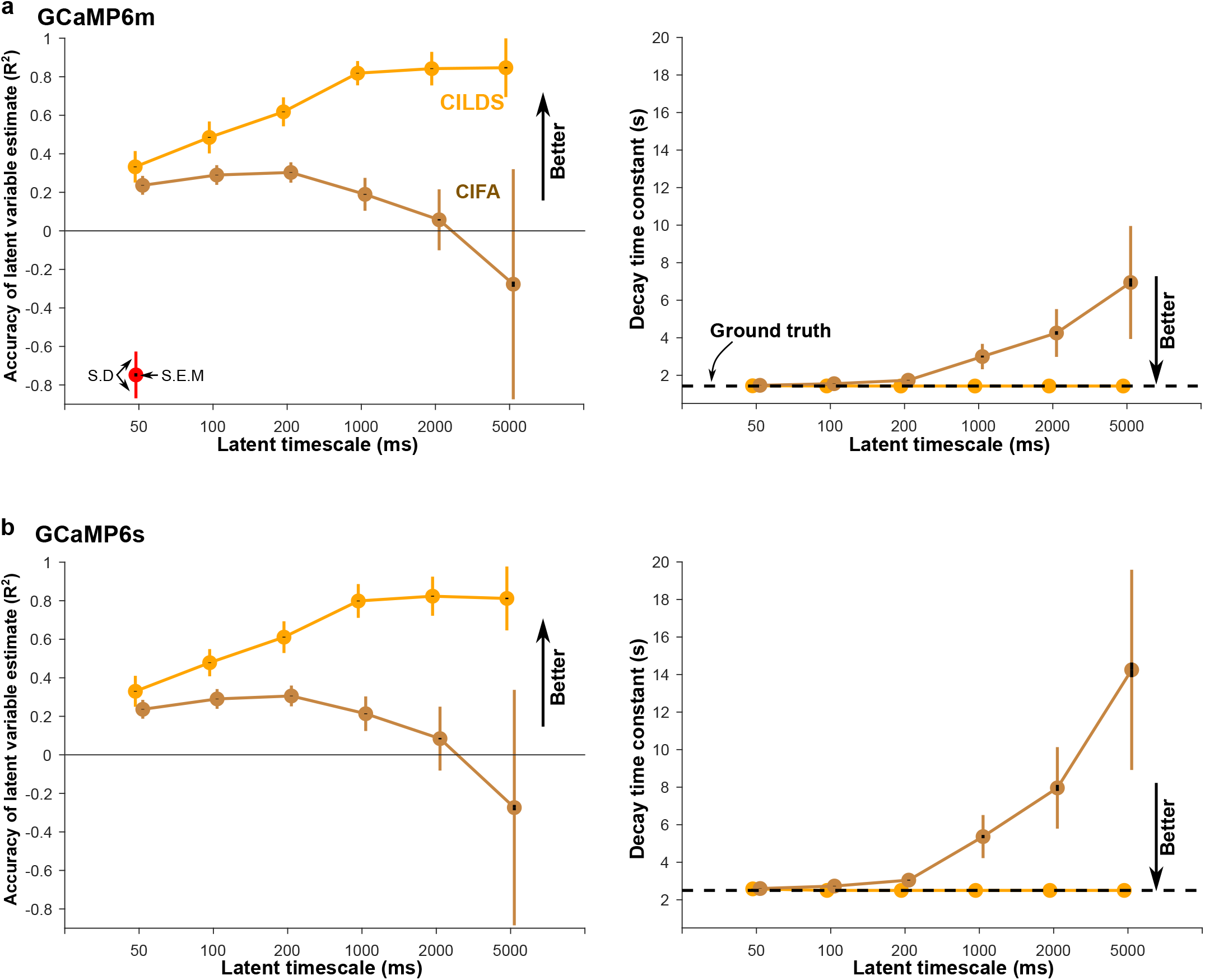
CILDS continues to outperform CIFA using different calcium indicators. Same conventions as Fig. 4. Here we compare between CILDS and CIFA for (a) GCaMP6m and (b) GCaMP6s, instead of GCaMP6f (Fig. 4). Consistent with Fig. 4, the overestimation of the calcium indicator decay time constant for CIFA increases as the latent timescales increase. The overestimation also increases when progressing from GCaMP6f (Fig. 4, note different vertical scale) to GCaMP6m (panel a) to GCaMP6s (panel b).

**Supplementary Figure 4.**
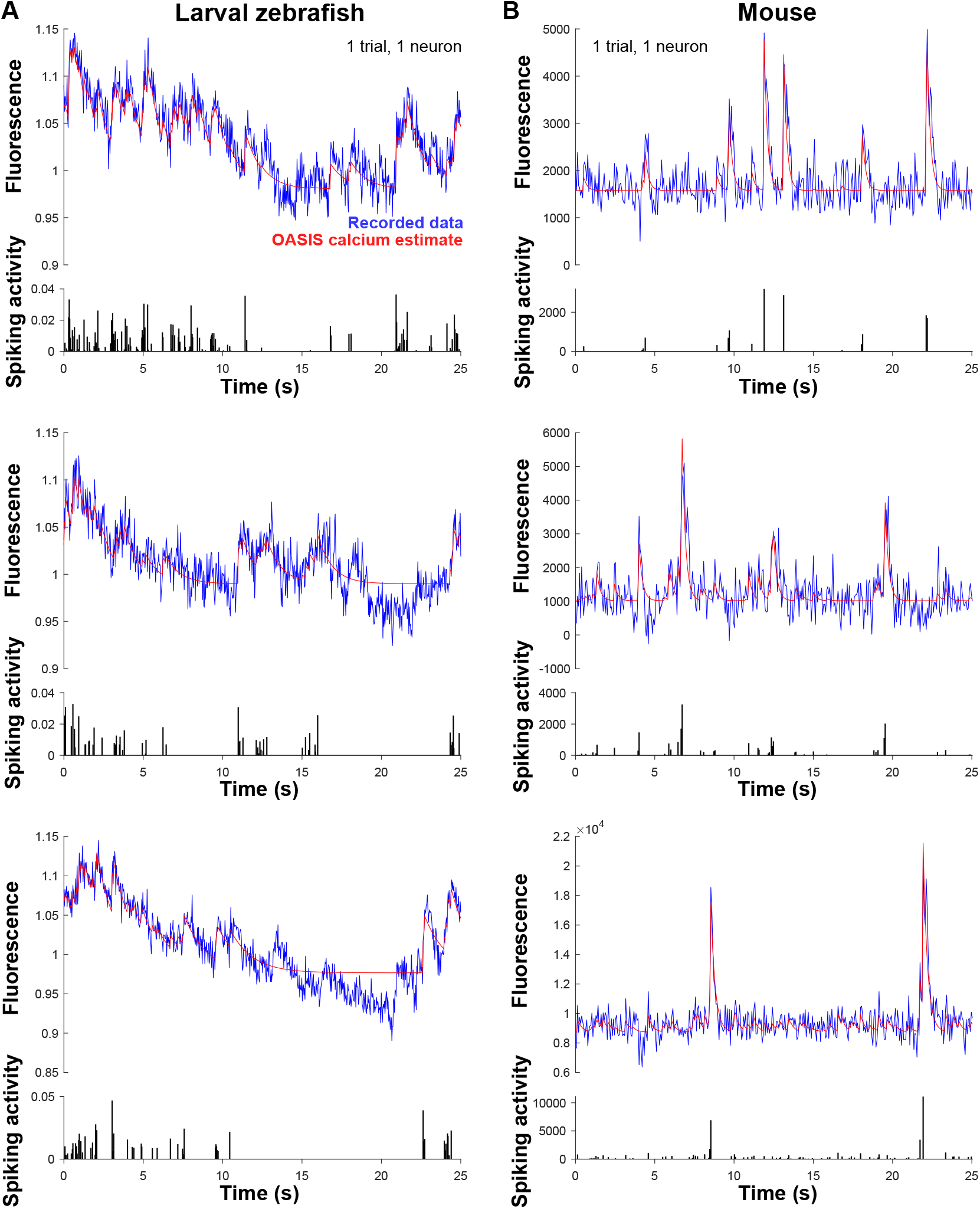
Examples of OASIS deconvolution, the first stage of deconv-LDS. **(a)** OASIS^34^ applied to recorded fluorescence traces (blue) to estimate the calcium time course (red) and spiking activity (black) for larval zebrafish. Each row shows 25 seconds of a different example neuron and trial from the same fish. **(b)** Same conventions as (a), but for recordings in mice. Note that no dimensionality reduction has been applied at this stage.

**Supplementary Figure 5.**
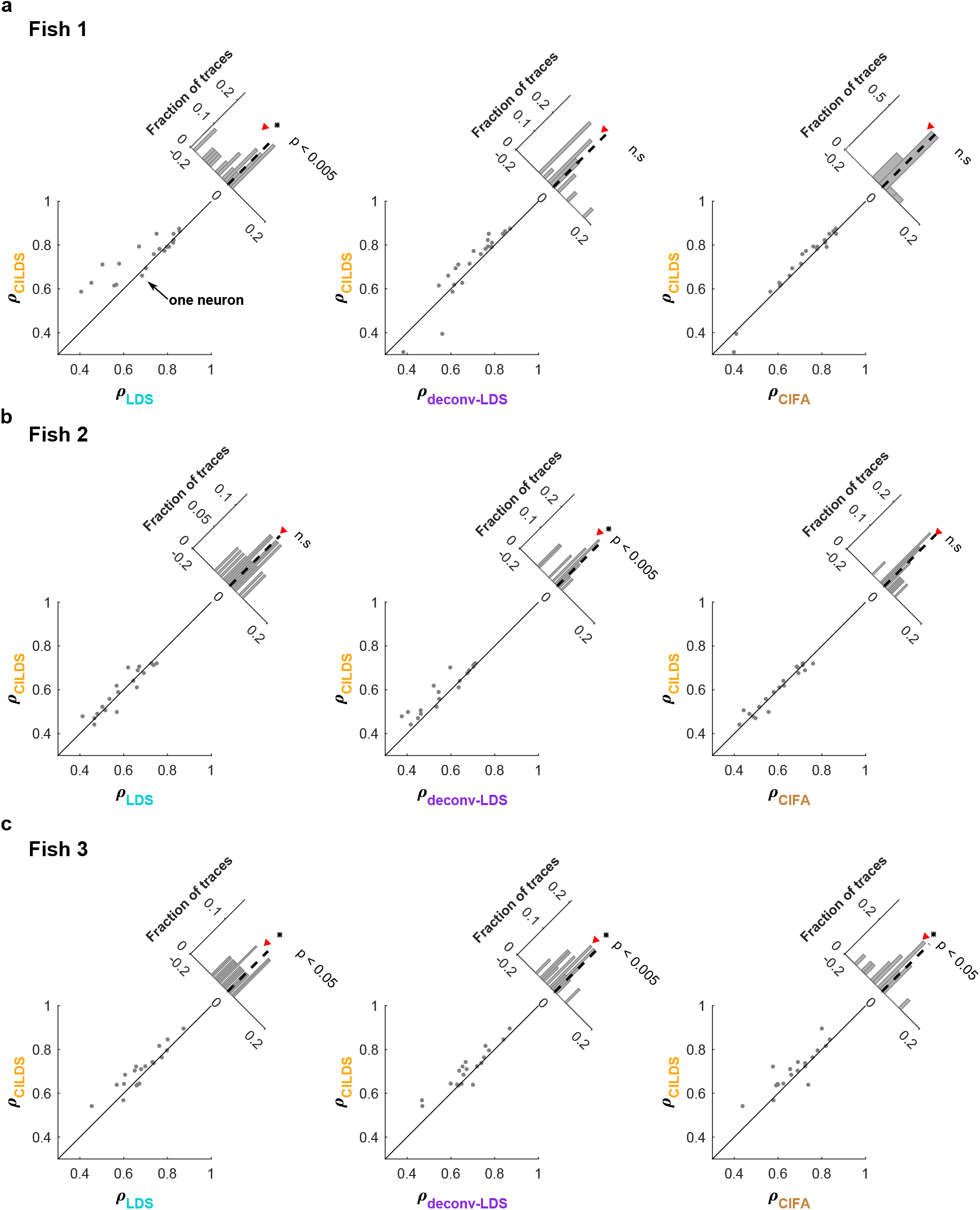
CILDS outperforms the other methods for individual larval zebrafish. Same conventions as Fig. 5c-e. **(a)** From left to right, correlations between recorded fluorescence and leave-neuron-out predicted fluorescence, comparing CILDS to LDS, CILDS to deconv-LDS, and CILDS to CIFA for Fish 1. **(b)** Same as (a), but for Fish 2. **(c)** Same as (a), but for Fish 3. These results are consistent with Fig. 5c-e, even at the level of individual fish. With an average of 20 points (i.e., neurons) per scatter plot, statistical significance is not attained in several plots, but in each case CILDS has a higher mean correlation than the other methods.

**Supplementary Figure 6.**
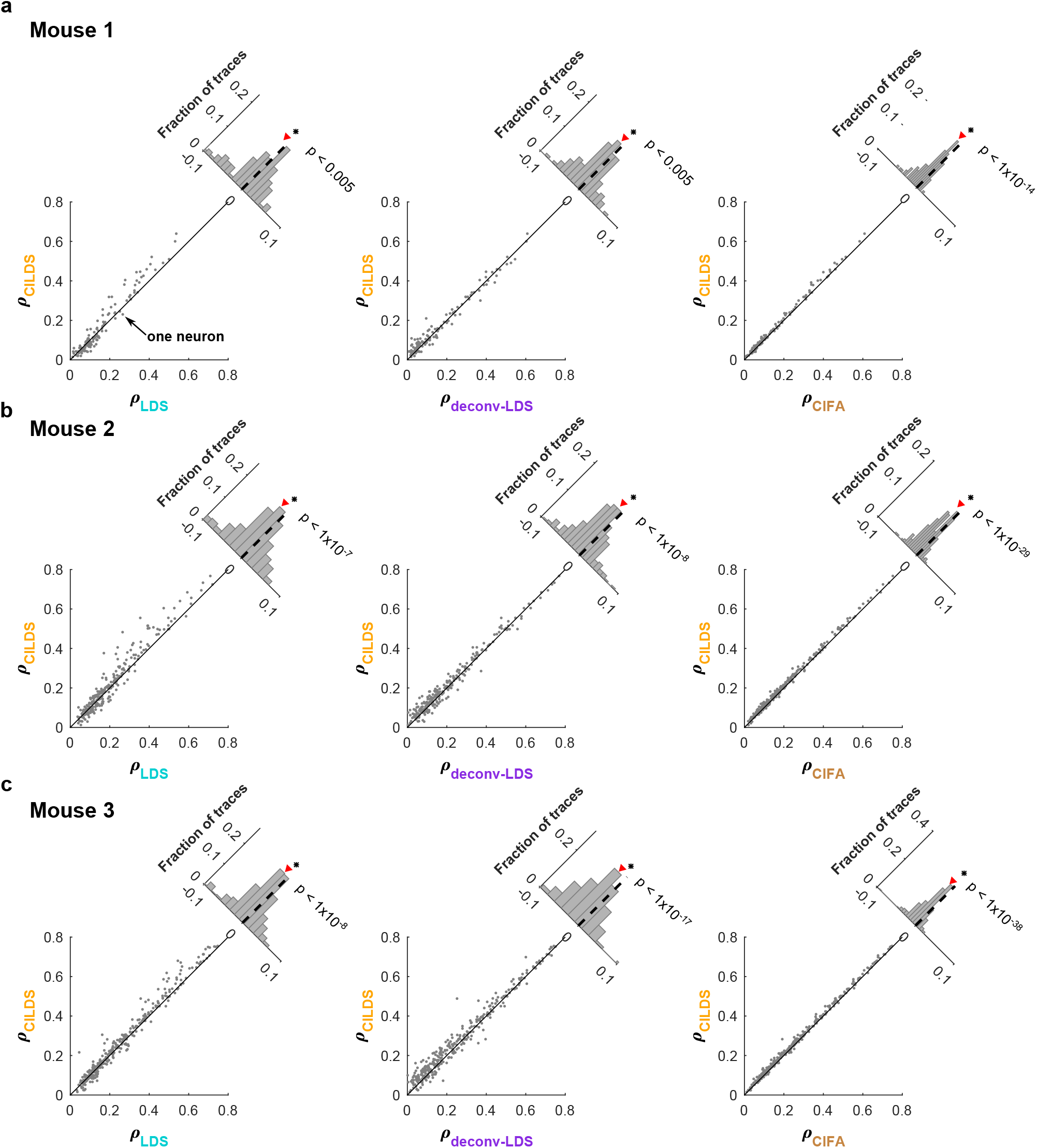
CILDS outperforms the other methods for individual mice. Same conventions as Fig. 6c-e. **(a)** From left to right, correlations between recorded fluorescence and leave-neuron-out predicted fluorescence, comparing CILDS to LDS, CILDS to deconv-LDS, and CILDS to CIFA for Mouse 1. **(b)** Same as (a), but for Mouse 2. **(c)** Same as (a), but for Mouse 3. These results are consistent with Fig. 6c-e, for each mouse individually.

**Supplementary Table 1.**
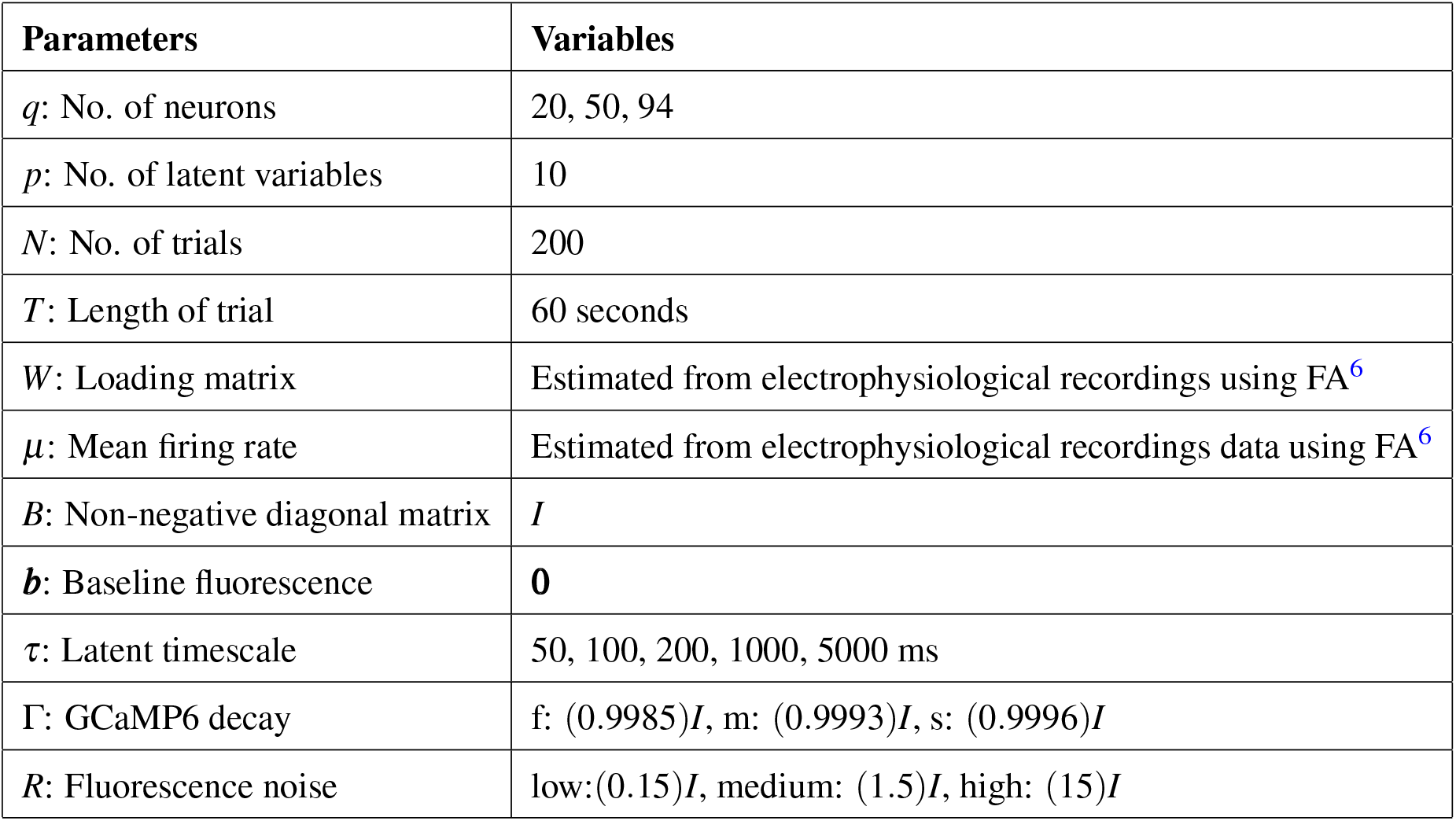
Parameter values used for simulations. Parameters within the simulation framework that were varied, and the range of parameter values tested. We chose parameter values to mimic those in real data.

## Supplemental Information

### EM algorithm for CILDS

The CILDS model is defined by equations (6) - (10). Only the fluorescence values ***y***_*t*_ are observed, whereas the calcium concentrations ***c***_*t*_ and latent variables ***z***_*t*_ are not observed.

The goal of the EM algorithm is to maximise the probability of the observed fluorescence traces *P*({***y***}) with respect to the model parameters *θ* := {*D, P*, ***h***_2_, *G*_2_, Γ, *A*, ***b***, *Q, B, R*, ***µ*_1_**,*V*_1_}, where {***y***} is shorthand for ***y***_1_, …, ***y***_*t*_. To perform this maximization, we iteratively perform an expectation step (E-step), then a maximization step (M-step), as detailed below.

#### 1 Expectation Step

The goal of the E-step is to compute the posterior distribution *s* := *P*({***c***}, {***z***}|{***y***}). Using this posterior distribution, we can compute the following expectations

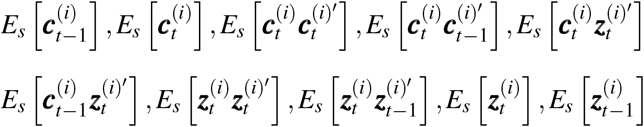

which are needed in the M-step. We start by rewriting equations (7) and (9) in block matrix notation

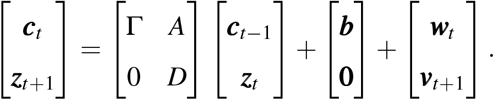

The observation model (6) can be written as

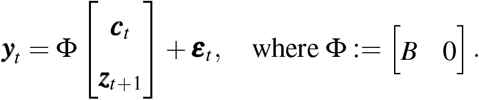

We define 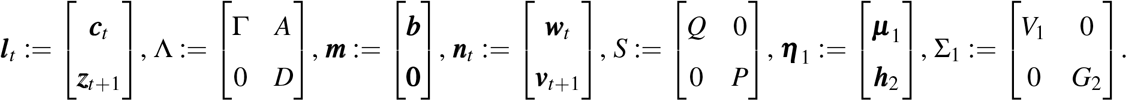

The CILDS model can thus be written in block matrix notation as

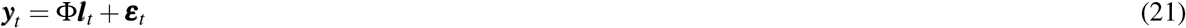

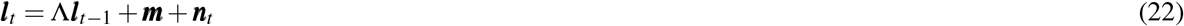

where ***l***_1_ ∼ ℕ(***η***_1_, Σ_1_), ***n***_*t*_ ∼ ℕ(**0**, *S*), and *t* = 1, …, *T*. In other words, CILDS can be written as an LDS whose parameters are constrained in a specific way. We seek to compute 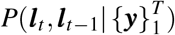 for *t* = 2, …, *T*. This distribution is Gaussian, and thus it is sufficient to find its mean and covariance. We denote 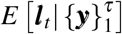 *b*y 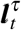 and 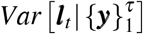 by 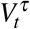, as in ref.^59^. To obtain the forward and backward recursion equations, we followed the steps outlined in ref.^64^. For brevity, we only show the results of the derivations below.

##### 1.1 Forward Recursions

To obtain 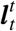 and 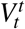, we recursively compute the following equations from *t* = 1 to *t* = *T*

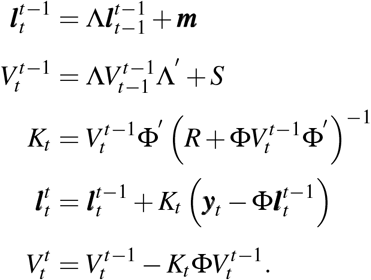

The recursions are initialized with 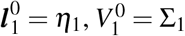.

##### 1.2 Backward Recursions

To obtain 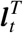 and 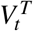, we recursively compute the following equations from *t* = *T* to *t* = 2. We also compute the covariance of the joint posterior distribution 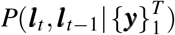, denoted as 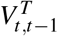

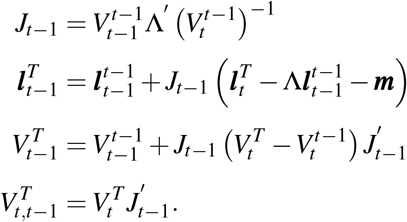

#### 2 Maximization Step

In the M-step, we seek to maximize the expected log joint distribution

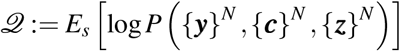

with respect to the model parameters, where *s* := *P*({***c***}^*N*^, {***z***}^*N*^ |{***y***}^*N*^ ; *θ*) and {}^*N*^ represents all *T* time points across all *N* trials. The joint distribution for one trial can be factorized as

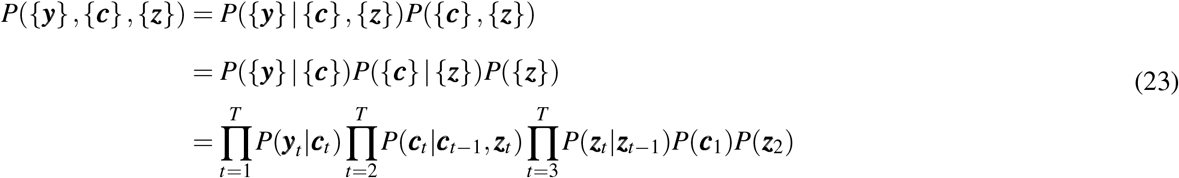

where these distributions are defined in equations (6) - (10).

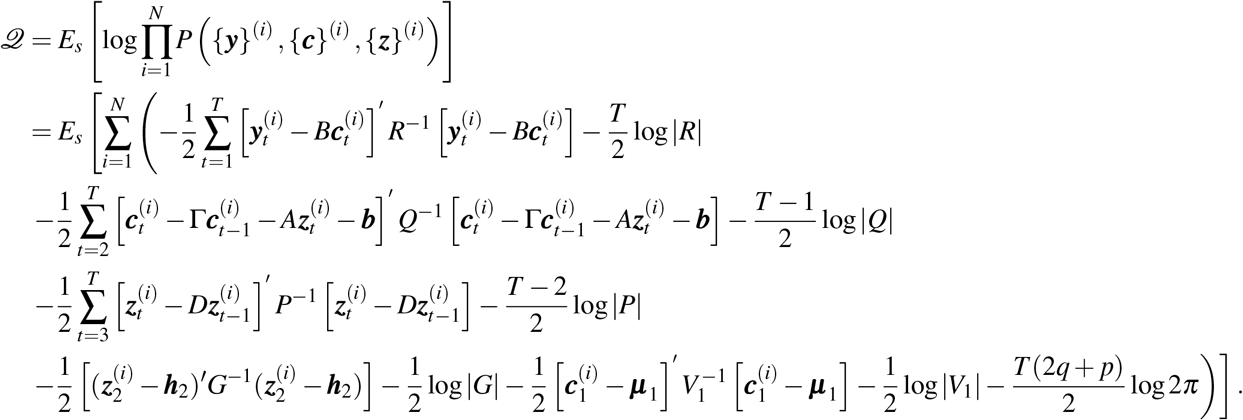

To maximize *𝒬* with respect to the model parameters *θ*, we compute the following partial derivatives

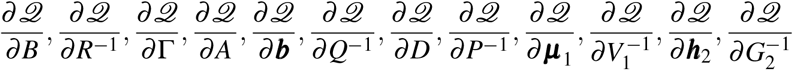

and set them to zero to solve for the parameters. Doing so results in the following M-step parameter updates, all of which can be expressed in closed form.

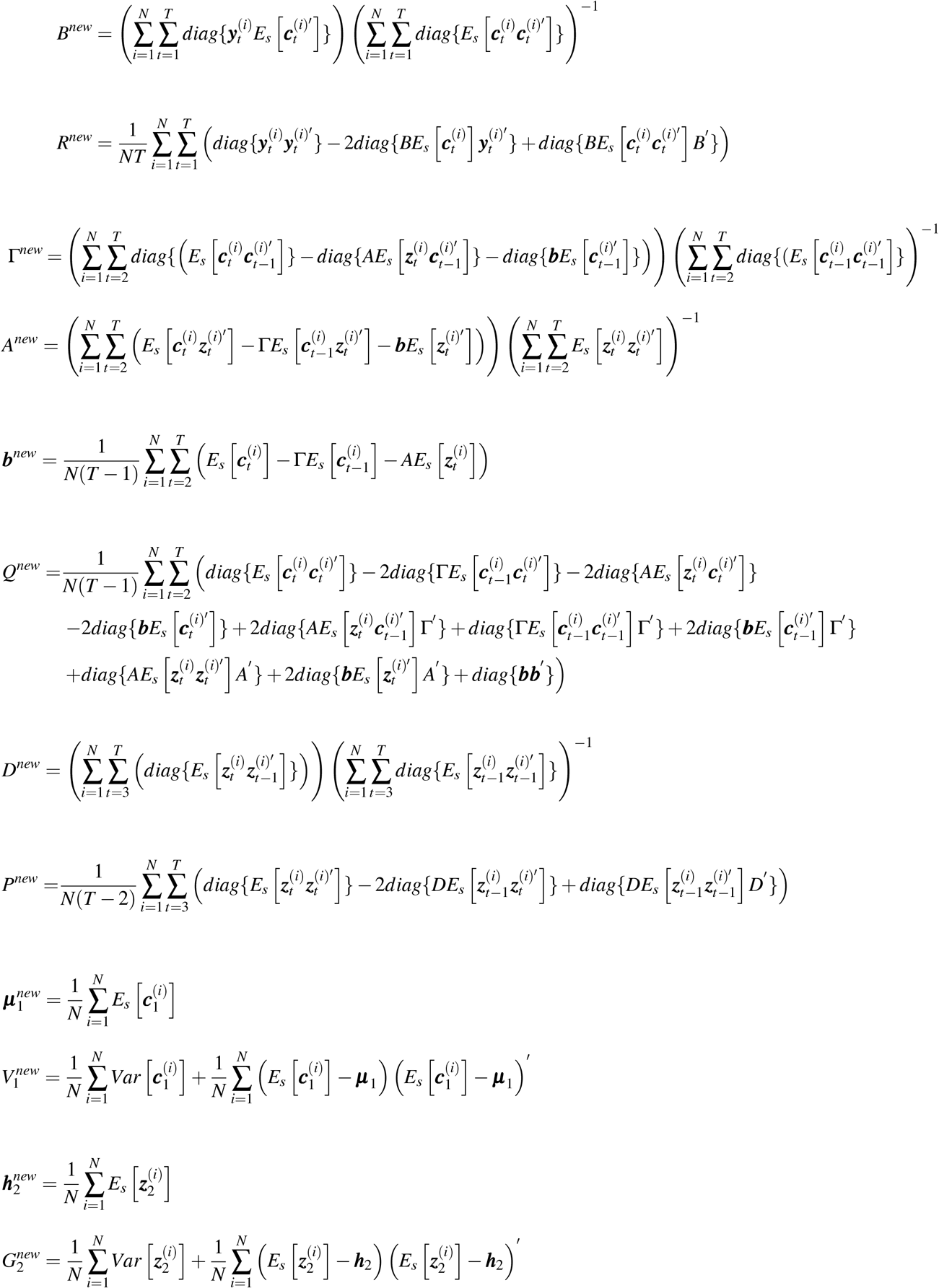

## Notes

### Competing Interest Statement

The authors have declared no competing interest.

